# Ghost interactions: revealing missing protein-ligand interactions using AlphaFold predictions

**DOI:** 10.1101/2023.10.18.561916

**Authors:** Nahuel Escobedo, Tadeo Saldaño, Juan Mac Donagh, Luciana Rodriguez Sawicki, Nicolas Palopoli, Sebastian Fernandez Alberti, Maria Silvina Fornasari, Gustavo Parisi

**Affiliations:** Departamento de Ciencia y Tecnología, Universidad Nacional de Quilmes, Bernal, Argentina

**Author notes:** Equally contributed (Nahuel Escobedo, Tadeo Saldaño).

## Abstract

Protein–ligand interactions represent an essential step in understanding molecular recognition, an intense field of research for many scientific areas. Structural biology has played a central role in unveiling protein-ligand interactions, but current techniques are still not able to reliably describe the interactions of ligands with highly flexible regions. In this work we explored the capacity of AlphaFold2 (AF2) to estimate the presence of interactions between ligands and residues belonging to disordered regions, which we called “ghost interactions” as they are missing in the crystallographic derived structures. We found that AF2 models are good predictors of regions associated with order-disorder transitions. Additionally, we found that AF2 predicts residues making ghost interactions with ligands, which are mostly buried and show a differential evolutionary conservation. Our findings could fuel current areas of research that consider intrinsically disordered proteins as potentially valuable targets for drug development, given their biological relevance and associated diseases.

## Introduction

Protein interactions are essential for sustaining life. Without exaggeration, Alan Fersht famously stated that “Biology is dominated by the chemistry of the noncovalent bond” ^1^. Noncovalent bonds are responsible for the folding of proteins into their functional shapes which defines and modulates the interaction pattern with their cognate ligands. As proteins show different degrees of conformational diversity ^2^, it is expected that those regions participating in ligand interaction tuned their flexibility during evolution to optimise ligand binding, interaction and catalysis ^3–6^. Following the ensemble nature of the native state ^7^, the prevalent vision to explain protein-protein and protein-ligand interactions is the redistribution of the conformational states (i.e. population shifts) in which unbound conformers with the conformation able to bind ligands are observed ^8,9^, although small induced fit contributions were also detected ^10^.

Structural biology has played a central role in unveiling protein-ligand interactions, mainly through the analysis of crystallisation derived data ^11^. Co-crystallisation and ligand soaking experiments enabled the analysis of nonbonded interactions between ligands and proteins, allowing the characterization of binding sites and pockets, ligand transit, its stabilisation and conformational preferences, among others ^12–14^. Most of these interactions are mediated by key residues that are mostly evolutionary conserved, reflecting their biological importance ^15^. Prediction of these conserved residues in a protein has been the cornerstone of a myriad of bioinformatic tools to predict the functional properties of proteins ^16^. Structure based methods derived from crystallographic information, however, have limitations. Highly redundant structural databases limit the scope of protein repertories to explore protein-ligand binding dynamics ^17^. Additionally, several proteins contain highly flexible regions, which appear as missing residues in protein structures derived from the application of crystallographic techniques ^18^. These regions have been associated with different types of “disorder”, with some residues that appear as missing in certain conformations but are shown as ordered in other conformers ^19–21^. These order-disorder regions are associated with several important biological processes such as ligand versatility, catalysis, allosterism and cooperativity ^2,22–24^. Missing regions near ligand binding sites impair our understanding of the role of stabilising forces due to the lack of structural information.

After the breakthrough of AlphaFold2 (AF2) ^25,26^ and its outstanding accuracy predicting protein 3D models ^27–31^, in this work we evaluated how well AF2 predicts missing regions that could maintain interactions with ligands. To this end, we used a collection of experimentally based protein conformers derived from the CoDNaS database ^32^. As mentioned before, in some cases it is expected that a missing region in one conformer will be ordered in another. Taking these ordered regions in given conformers as a criterion of truth, we can estimate the AF2 modelling capacity of these missing regions (order-disorder regions or ODRs) using structural comparisons. We found that AF2 can predict missing regions (from 5 to 100 residues long) with high precision. Additionally, taking a large dataset of holo forms of proteins, we found that AF2 models predict segments of the structure that can potentially interact with ligands that are missing in the crystallographic structures due to their high flexibility or disorder. Furthermore, residues involved in these interactions, which we called “ghost interactions” as they are missing in crystallographic structures, are differentially evolutionary conserved and mostly buried, a finding that could indicate their biological relevance and predictability. Our results suggest that AF2 models could help in the study of protein-ligand interactions when ligands bind to binding sites surrounded by highly flexible regions.

## Results

### AF2 predicts order-disorder transition regions

Using CoDNaS ^32^ and MobiDB ^33^ we selected 5,477 proteins with different extensions of conformational diversity, measured by their structural differences between conformers quantified by their alpha carbon-based Root Mean Square Deviation (RMSD). These conformers are derived from experimental methods, commonly X-Ray crystallography. All the proteins selected have at least one conformer with at least one disordered region with more than 5 consecutive missing residues which is not missing in another conformer of the same protein (see Figure 1a). We call these regions “ODR” (order-disorder regions). In this dataset, each protein has on average 8 conformers and their disordered regions have lengths from 5 to 100 residues (longer lengths were removed from the dataset due to the low number of cases, see Figure 1c). The distribution of the number of missing residues per region of a protein is shown in Figure 1c.

**Figure 1:**
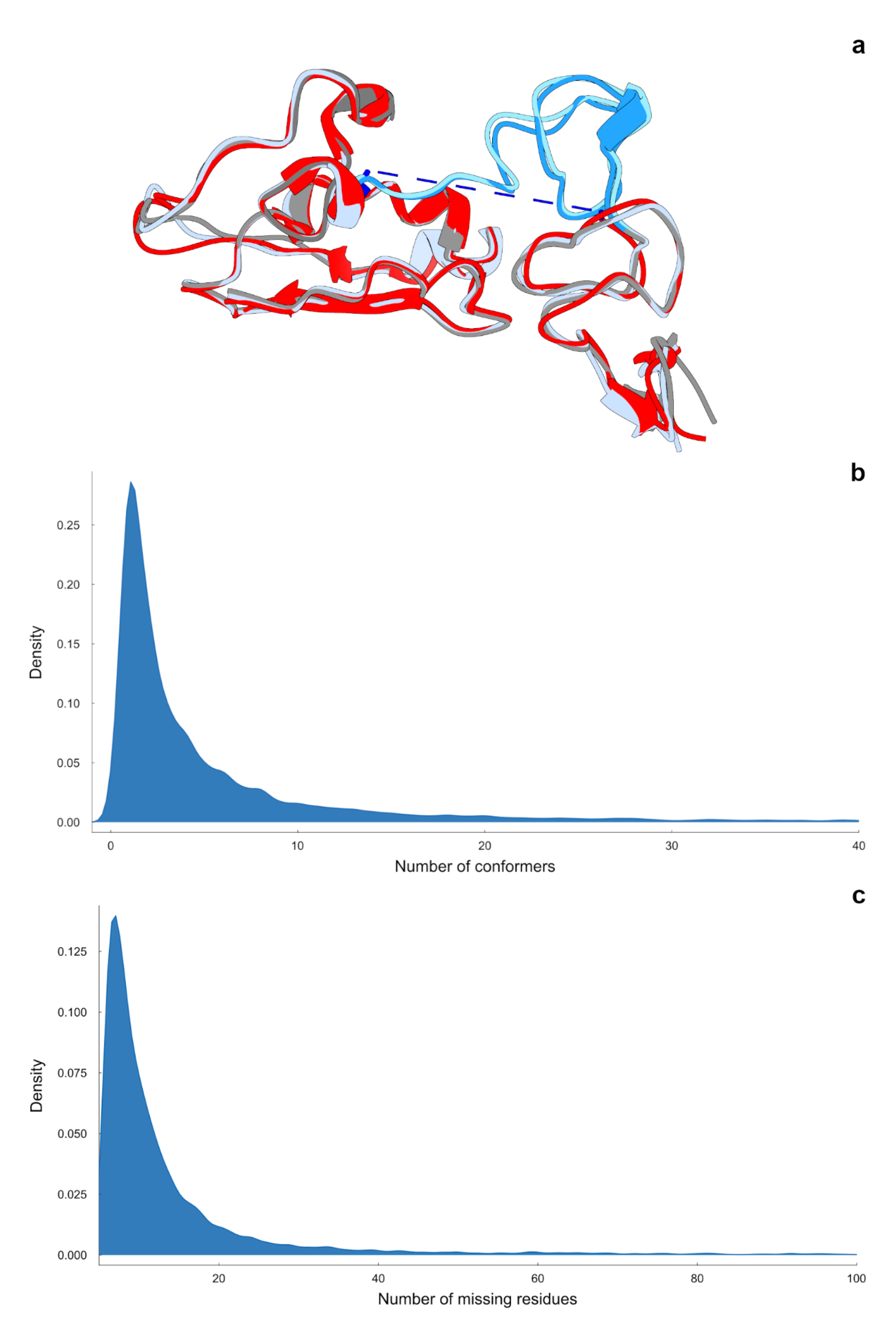
Schematic representation of the dataset. **a**. Three conformers from protein P00533 (PDB codes 4KRO in red, 3NJP in grey and 1NQL in light steel blue; structures are shown from T71 to R228 in PDB numeration). Structure 4KRO has a missing region between residues C183 to C208 (blue dashes), also predicted as a disordered region using MobiDB. However, the structures of two conformers (PDB 3NJP and 1NQL) show this region as ordered, represented in light blue in both structures. Consequently, this is marked as an order-disorder region or ODR **b**. Distribution of the number of conformers per protein in the dataset. **c**. Distribution of the length of missing regions per protein in the dataset.

Using the AF2 model and removing proteins with an average pLDDT score < 70, we retain 4,097 proteins with an average pLDDT =87.5 and 8 conformers on average per protein. With this final dataset we estimated the accuracy of AF2 on the prediction of ODRs. To this end, we structurally aligned the AF2 fragment corresponding to the missing region found in at least one of the conformers, to the structured region found in another conformer of the same protein (illustrated in Figure 1a, panel a). The resulting average RMSD for these ODR alignments is 1.16 Å (Figure 2). Splitting the dataset of disordered regions in half and according to their average pLDDT (below and above 70 on average) we observed that the average values of the RMSD distribution are 2.29 Å and 0.91 Å for the disordered regions below and above 70 pLDDT in average respectively (Figure 2, panel b). These results suggest that a pLDDT as low as 70 for the segment corresponding to the missing region is still a good predictor of the conformation a disordered region can adopt.

**Figure 2:**
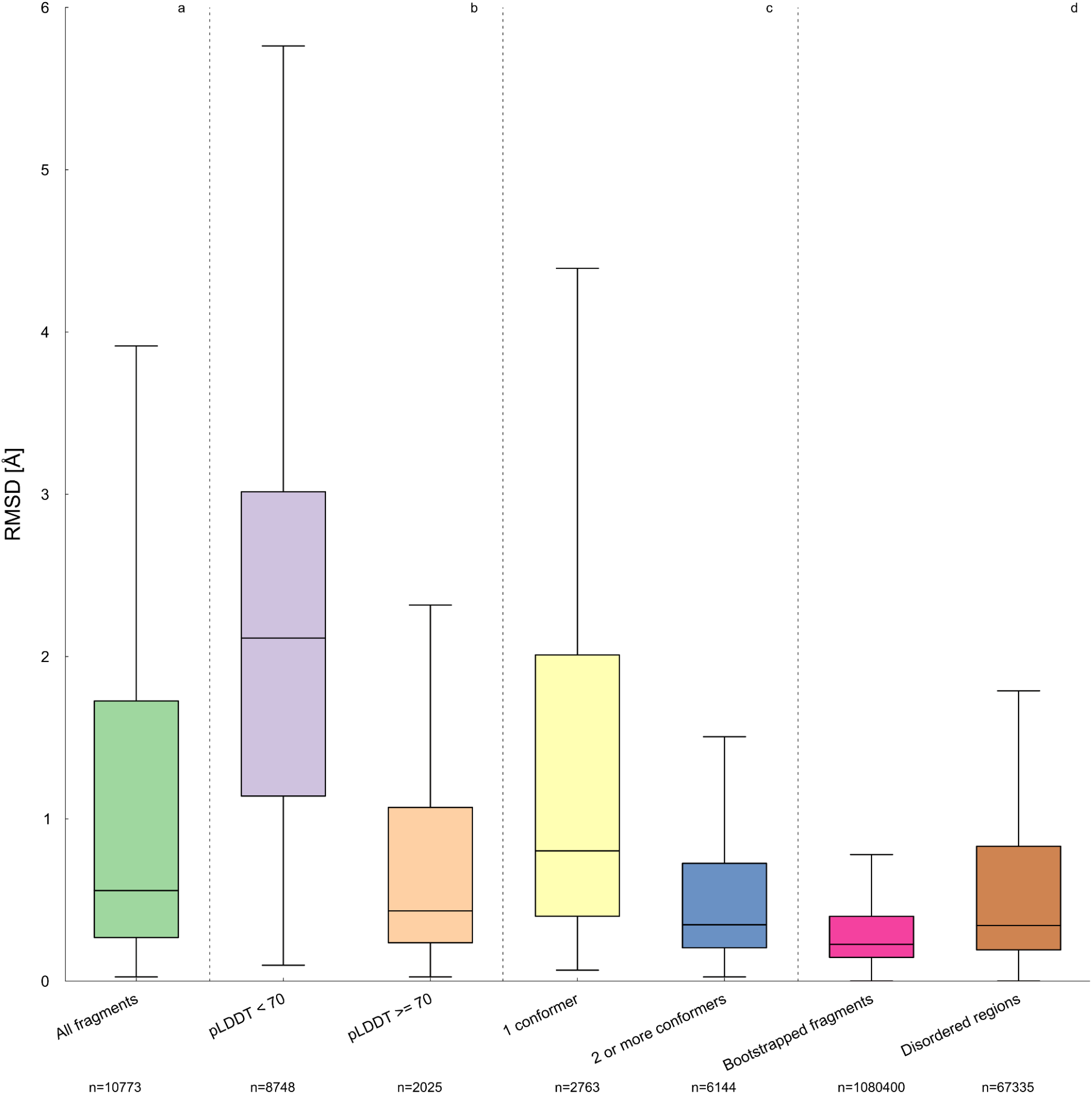
Structural assessment of AF2 capacity to predict order-disorder regions. RMSD distributions of segments of AF2 models aligned with respect to experimental conformers showing missing regions as ordered (panel a, green boxplot). Models with good predictive values (pLDDT >= 70) (panel b, orange boxplot) showed lower RMSD than models with poor pLDDT (panel b, purple boxplot) indicating that pLDDT is a good estimator of highly flexible conformations that can alternatively adopt ordered conformations. For those proteins with a disordered region that is ordered in two or more alternative conformers (panel c, blue boxplot), AF2 is able to better reproduce one of those ordered conformations, as revealed by the lower RMSD compared with those proteins with only one ordered conformation (panel c, yellow boxplot). Distribution of AF2 bootstrapped fragments corresponding to ordered regions (panel d magenta) and conformational diversity of ODRs (panel d, brown) are included as references. All the distributions in the same panel showed a Wilcoxon rank-sum test p-value < 0.01 between them, except Disordered regions and Bootstrapped fragments. Outlier points are not displayed here. (All fragments *n* = 10,773; pLDDT < 70 *n* = 2,025: pLDDT >= 70 *n* = 8,748; 1 conformer *n* = 2,763; 2 or more conformers *n* = 6,144; Bootstrapped fragments *n* = 1,080,400; Disordered regions *n* = 67335).

In 2,747 proteins with pLDDT > 70, with 6,052 fragments in total, we found more than two conformers showing the ODR as ordered. Recording the best fit of the AF2 fragments with the different ordered fragments coming from different conformers we found a slightly lower RMSD (0.71 Å, Figure 2, panel c). For the rest of the proteins that only have one conformer with the ordered ODR, the RMSD between those segments and AF2 gives an average RMSD of 1.36 Å (Figure 2, panel c). To put these results into perspective, we compared these distributions with the error rate of AF2 when predicting ordered regions. To this end, for each evaluated protein we applied a bootstrapping protocol, in which we structurally compared an ODR with each of 100 resampled fragments of the same length of the corresponding ODR, taken from the same protein. This distribution is shown in Figure 2, panel d, and has an average RMSD = 0.50 Å. Additionally, we estimated the conformational diversity of the ODR regions using the set of proteins with more than one conformer showing the ODR as ordered (Figure 2, panel d). Recording the maximum RMSD between those fragments as a proxy to the conformational diversity of the region, it is observed that the distribution of these flexible fragments is RMSD = 0.86 Å on average. As a result of these analyses we observed that AF2 is able to predict ODRs with high confidence in the best of the analysed cases with an average RMSD = 0.71 Å when conformational diversity is considered. Interestingly, we found that this distribution is in the same order as the conformational diversity of the flexible region of the ODR (Wilcoxon rank-sum test p-value > 0.01).

As missing regions in our dataset have different lengths, we also explored the impact of the ODR length in AF2 predictions. We found that AF2 performance is almost independent of the ODR length. Figure 3 shows the RMSD distributions as a function of ODR length bins. Averages in RMSD in each length bin comprise similar average values and interquartile ranges, with no clear dependence on ODR length (1.11, 1.41, 1.64, 1.50 and 1.19 Å respectively). See also Supplementary Figure 1.

**Figure 3:**
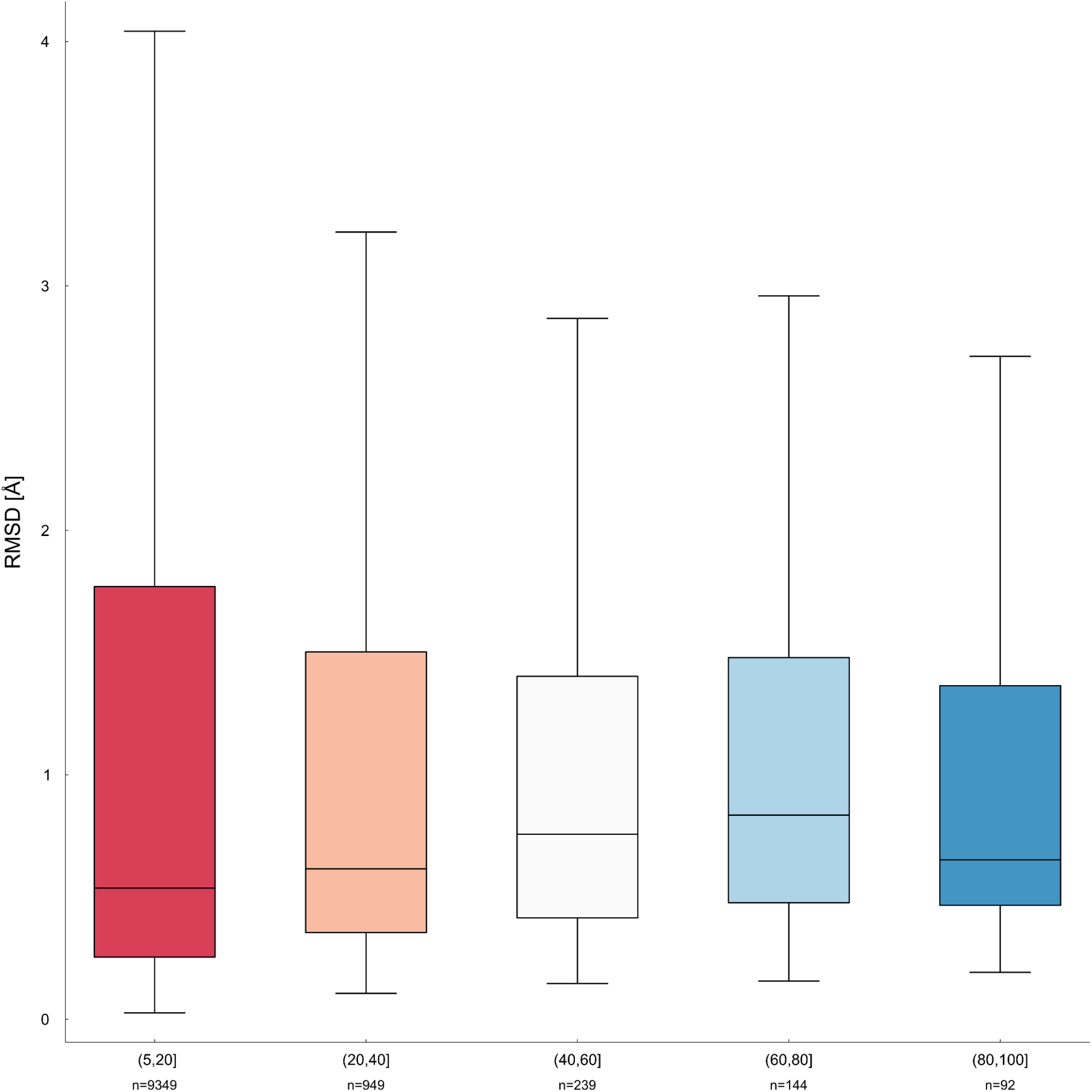
RMSD distributions of ODRs and AF2 models as a function of ODR lengths. The number of proteins in each bin are as follows: (5, 20] *n* = 9349, (20, 40] *n* = 949, (40, 60] *n* = 239, (60, 80] *n* = 144, (80, 100] *n* = 92). For all Wilcoxon rank-sum test, see Supplementary Table 2.

### Predicting contacts with ligands using AF2

When protein structures are co-crystallised with cognate ligands, the resulting data can be used to characterise protein-ligand interactions associated with binding and/or with catalytic processes, revealing intrinsic mechanistic explanations of protein biology. To evaluate the capacity of AF2 to predict ligand-protein interactions, we selected 11,323 proteins (55,773 PDB structures) with cognate ligands, thus representing holo forms of the proteins from PDB (see Methods). AF2 models for those proteins showed an average pLDDT of 87.4 and a global RMSD with each corresponding holo form of 1.91 Å, showing that AF2 models are of good quality (see Supplementary Figure 2 for pLDDT and RMSD distributions). The experimental structures of the holo forms were structurally aligned with their corresponding AF2 models, the cognate ligands were “transplanted” into the AF2 models, and contacts between each AF2 model and the transplanted ligand were then estimated (see Methods). We also explored the quality of the AF2 protein-ligand interactions, estimating the clashes using a measure called the “transplant clash score” (TCS) derived previously ^34^. When TCS was estimated in the holo forms captured in PDB files and in the AF2-ligand interactions obtained after structural alignment, the distributions of clashes overlapped (see Supplementary Figure 3).

Using the AF2 models and the holo forms, we found that AF2 predicts contacts with a median relative error of ∼8% (Supplementary Figure 4, 5 and Figure 4), defined as the difference between the number of PDB contacts and the number of AF2 model contacts, normalised by the number of PDB contacts. This relatively low error also indicates that aligning the holo form with the AF2 model resulted in relatively good positioning of the ligand in the AF2 models. As shown in Figure 4, AF2 models conserved the great majority of the contacts found in ordered regions of the PDB structure although some structures show variations above or below the PDB contacts.

**Figure 4:**
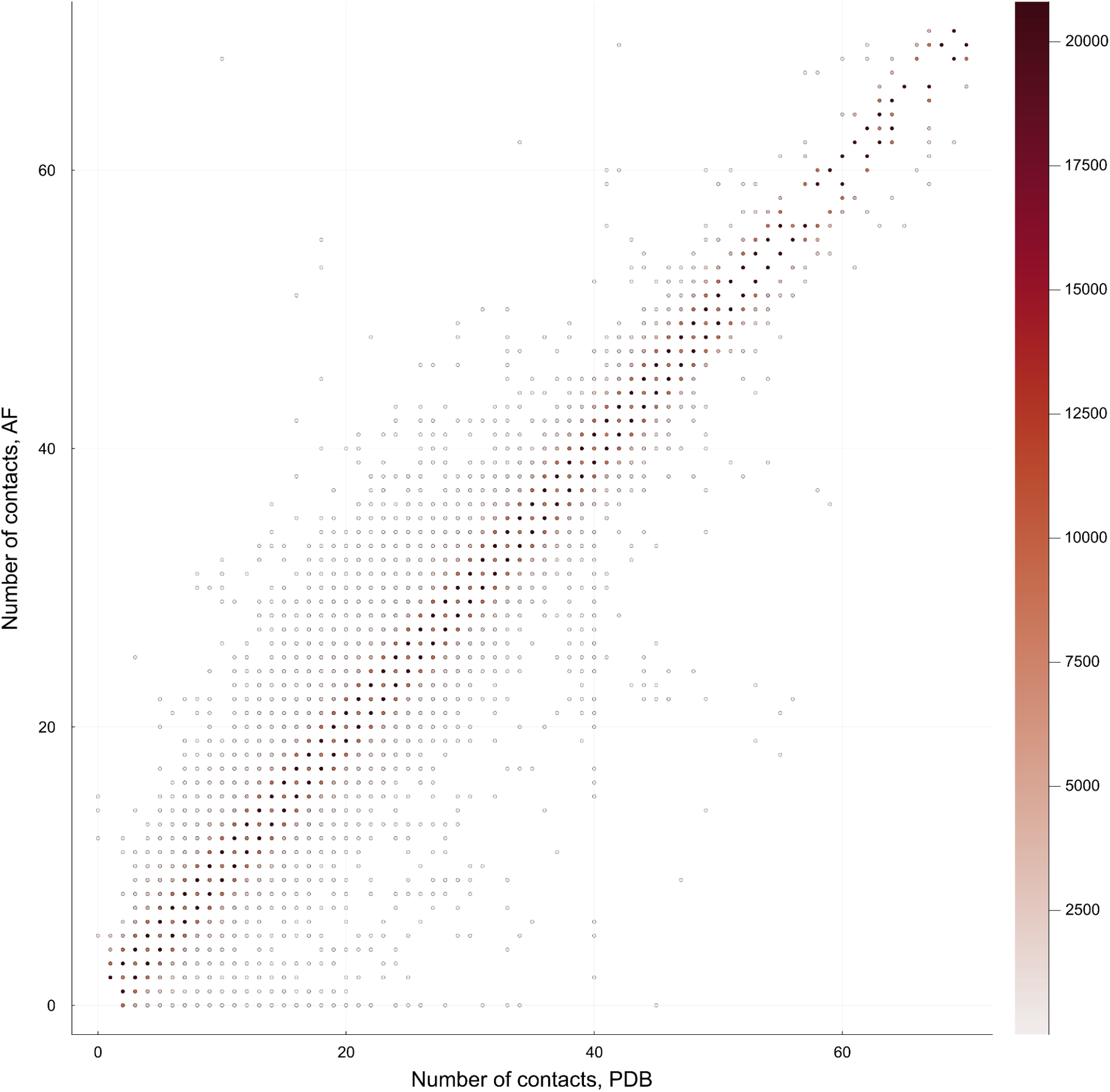
Comparison of the number of contacts of both PDB and AF2 structures with the ligand. Point darkness indicates the density of structures with a given number of contacts in the PDB holo form and in the AF2 model, according to the scale on the right. Points near and on the line y=x indicate that AF2 accurately reproduces the number of contacts with ligands found in PDB holo forms (Pearson correlation ρ = 0.94, p-value < 0.01).

We also found that the relative error shows a weak correlation with the global pLDDT model (Pearson correlation ρ = −0.177104, p-value < 0.01) and with the RMSD between the model and the holo structure (Pearson correlation, ρ = 0.337735, p-value < 0.01). According to these correlations, structural deviations measured with RMSD are more informative for detecting deviations in the number of contacts than pLDDT. Finally, we also found that the relative error is independent of the molecular weight of the ligand (Pearson correlation ρ = −0.08, p-value < 0.01).

### Detecting “ghost” interactions revealed by AF2 predictions

Given that we have found that AF2 is able to predict conformations of disordered regions that undergo order-disorder transitions (ODRs), we are interested in the evaluation of AF2’s capability to predict non-covalent interactions between disordered regions and ligands. Disordered regions are mostly associated with the presence of missing regions in proteins ^19,20^. When these regions are missing due to their intrinsic flexibility, essential information about those interactions is also lost.

Using the protein-ligand contact information and the reported missing regions (disordered regions) in the corresponding holo forms, we filtered the dataset to keep only those proteins showing AF2 contacts with the ligand in regions that are missing in the PDB files (456 proteins, 1,113 PDB structures). These are called “ghost interactions” from now on. These AF2 regions showed a relatively good average pLDDT = 73.82 (Supplementary Figure 6, panel a) despite the fact that they are likely very flexible or disordered. The distribution of the lengths of missing regions containing residues with ghost interactions has an average value of 13.59 amino acids (Supplementary Figure 6, panel b; also see Supplementary Figure 7 for some examples). With this length distribution and according to the results in Figure 3, these regions have an average error measured by the RMSD of 1.11 Å. As a result we obtained an average of 3.31 residues making ghost interactions per structure.

Can we rely on the contacts between disordered regions modelled by AF2 and cognate ligands? We hypothesise that residues responsible for interactions with cognate ligands should be evolutionary conserved in order to sustain the biological functional activity. To test the quality of the predicted interactions of residues showing ghost interactions, we estimated their evolutionary conservation compared with observed protein-ligand contacts belonging to ordered regions in the PDB. These results are shown in the left panel of Figure 5. Residues belonging to ghost interactions are as much evolutionary conserved as are the residues with contacts belonging to ordered regions. Furthermore, these residues are significantly more conserved than flanking residues in the same missing region that do not present ghost interactions (the average values for residues with contacts with the ligand in ordered regions, residues with ghost interactions, and missing residues are 0.64, 0.56, and 0,48, respectively). The same tendency was observed when we analysed the pLDDT values for the three groups (average values of 90.09 for residues with contacts in ordered regions, 75.05 for residues with ghost interactions and 71.30 for missing residues) and the Relative Solvent Accessible Area, RSA (average values of 0.13, 0.21 and 0.37). The three measures reveal equivalent tendencies, showing that residues establishing “ghost interactions’’ are differentially conserved and buried with a similar behaviour to residues in ordered regions of the protein making contacts with ligands.

**Figure 5:**
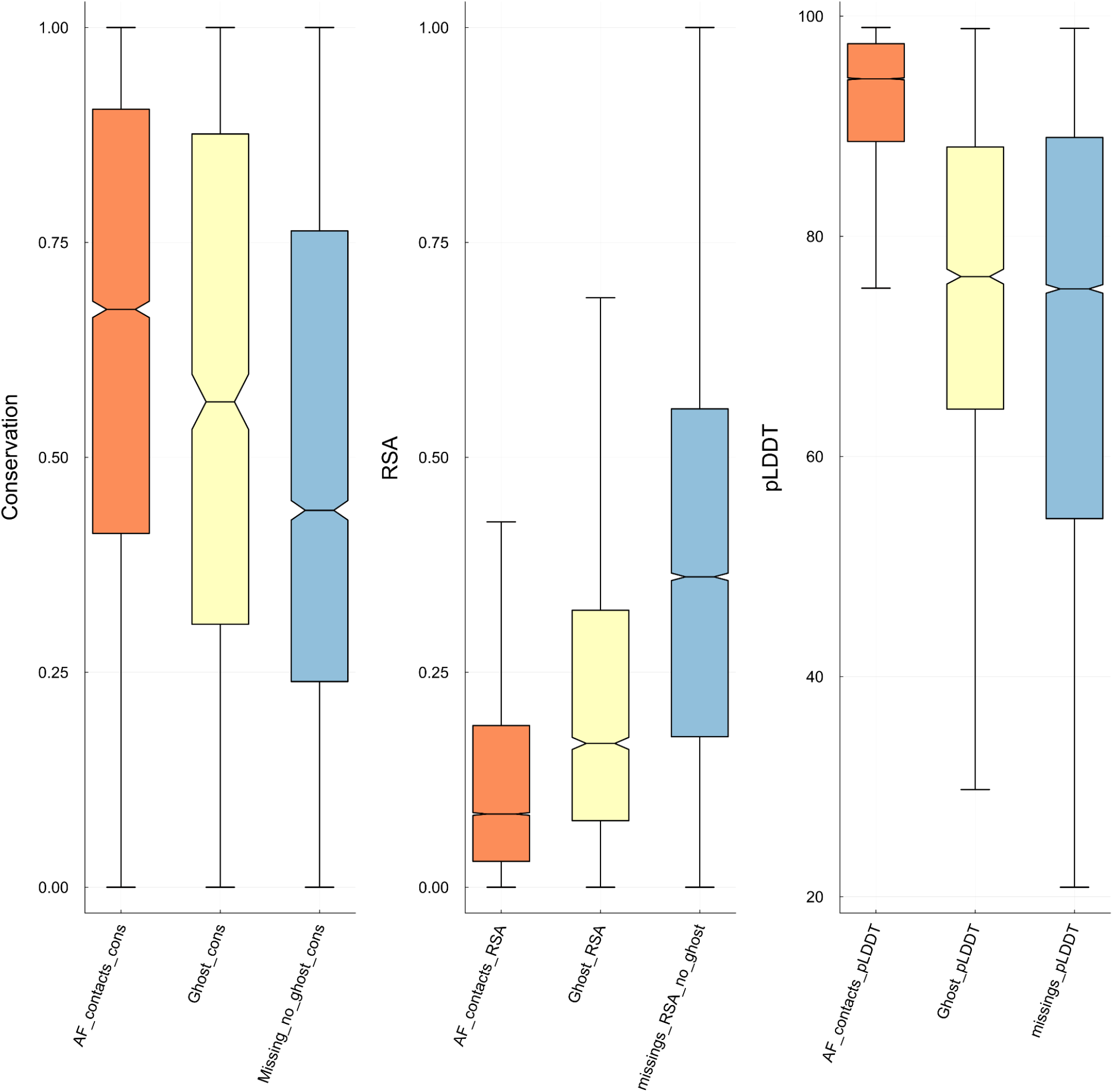
Evolutionary conservation, RSA and pLDDT of different evaluated positions. Here we compared three different sets of residues: residues in missing regions (light blue), residues in ordered regions that make contacts with ligands (orange), and “ghost” (light yellow) residues. Outlier points are not displayed. For all values, the Wilcoxon rank-sum test p-value between them is p-value < 0.01.

We found significant differences between the amino acid distribution of residues making ghost interactions compared with the distribution of residues in ordered regions that make contacts with ligands (See Supplementary Figure 8). We observed that ghost interactions are enriched in Arg, Gln, Ser, Gly and Ala, while being depleted in Val, Met, Ile, Cys, Asn and Asp (see Supplementary Figures 8 and 9 and Supplementary Table 4). Although residues making ghost interactions are located in disordered regions, their amino acid distribution is completely different when compared with amino acid distribution of disordered residues coming from the Disprot database (See Supplementary Figure 9). These results indicate the uniqueness of residues making ghost interactions with respect to other residues commonly found in ordered or in disordered regions.

### Predictive Power of “ghost interactions”

To assess the capacity of AF2 to predict ghost interactions, we used Receiver Operating Characteristic (ROC) curves to assess the ability of various AF2-based metrics to predict ghost interactions with ligands. Given an ODR predicted as before, we determined if pLDDT, evolutionary conservation or relative solvent accessibility (RSA) were able to highlight residues within this region that may establish ghost interactions. Figure 6 shows the performance of these three measures in predicting ghost interactions from AF2 models, both calculated per residue as well as their averages derived from the corresponding fragments.

**Figure 6:**
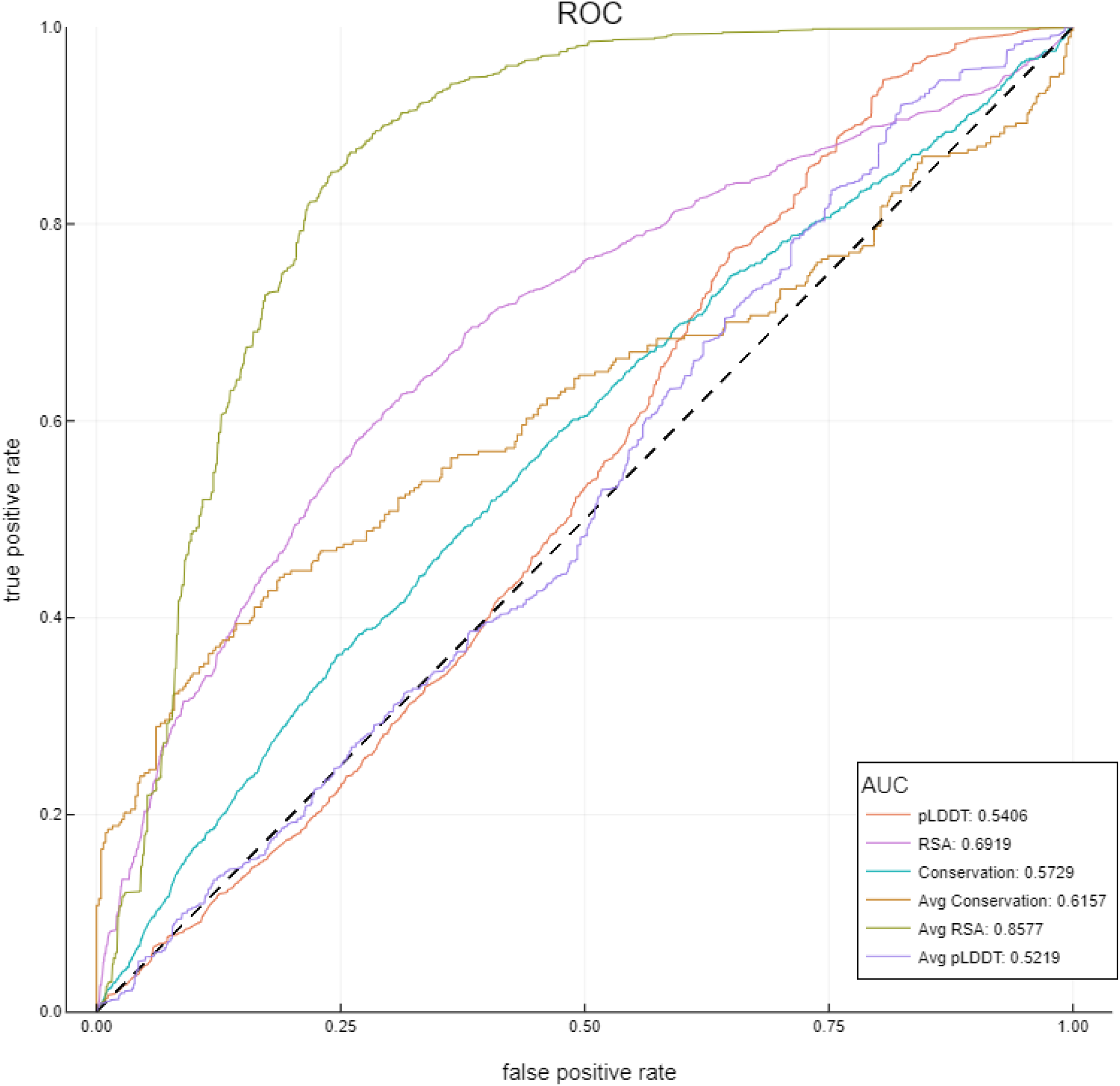
Predictive capacity of ghost interactions. Among different measures assayed, RSA (relative solvent accessibility) and average RSA for each region containing putative ghost interactions are the best predictive measures. RSA (relative solvent accessibility; pink line), pLDDT (orange line), Conservation (using unweighted variance measure; blue line), Average RSA (green line), Average pLDDT (purple line) and Average Conservation (brown line) are calculated as the average value of the respective measure for each PDB region.

According to this analysis, the best predictive measure is the average RSA estimated for the ODRs containing the residues that make putative ghost interactions (AUC = 0.85), followed by the plain RSA (AUC = 0.69). The two other parameters, evolutionary conservation and pLDDT, showed lower predictive capacity (AUC = 0.54 and 0.52, respectively) that only slightly improved by taking the average in each region (AUC = 0.57 and 0.61). Although residues making ghost interactions are more conserved and have a higher pLDDT than the rest of the residues in the missing region, this difference appears too small to reflect a good predictive capacity by itself. RSA, on the other hand, is by far the best predictor, distinguishing between “ghost” residues and the rest of the disordered positions in an ODR.

### Relevance of ghost interactions in biology

Human glucokinase (GK), an enzyme that acts as a main regulating sensor of plasma glucose levels, is a clear example of the relevance of the information provided by ghost interactions. Inactivating mutations in GK are associated with diabetes, while activating mutations are linked to hypoglycemia. GK catalyses the conversion of glucose to glucose 6-phosphate ^35^. Unlike other hexokinases, GK exhibits lower glucose affinity and positive cooperativity, and is not affected by product inhibition ^36,37,38^. Glucokinase activators (GKAs) stabilise the allosteric site, promoting active conformations. Due to the critical role of GK in glucose homeostasis, several GKAs have progressed to clinical studies. These GKAs show promise in effectively treating hyperglycemia but occasionally lead to hypoglycemia as a significant risk. In efforts to mitigate the risk of hypoglycemia while maintaining efficacy, ‘partial activators’ of glucokinase have been investigated. 2-Methylbenzofurans derivatives have been identified as promising candidates ^39^. The GK structure in complex with one of these ligands (PDB: 3S41) reveals several interactions, including a dimethyl pyrimidine carboxamide interacting with the side chain of Y215, residue in helix C, and V91 and V101 from the disordered loop 91–101. The AF2 structure allows us to observe a ghost interaction of W99 with this ligand (Figure 7a) that contributes to the stabilisation of the ligand in the binding site and has the potential to influence the observed active conformation. Interestingly, this key position was previously identified as the site of a novel activating GK mutation (p.Trp99Arg) associated with familial hyperinsulinemic hypoglycemia ^40^. Understanding the activation mechanism of GK and the interactions involved in it will aid in the design of GKAs with improved safety profiles for the treatment of conditions arising from activating mutations.

**Figure 7:**
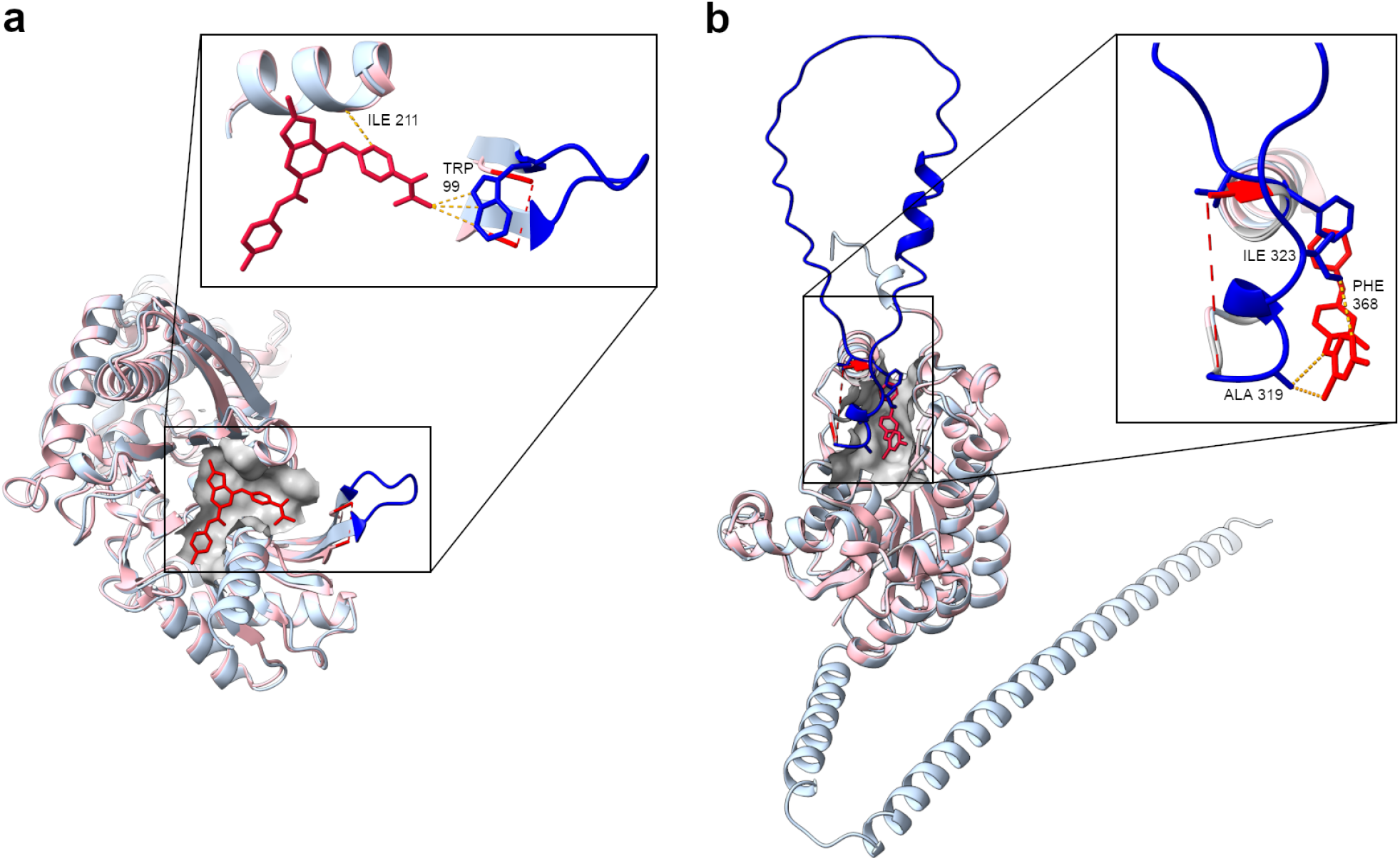
Glucokinase (GK) and *Plasmodium falciparum* enoyl acyl carrier protein reductase (PfENR) ghost interactions. GK (PDB: 3S41; panel a) and PfENR (PDB: 2OP1; panel b) are shown in pink. AF models of each protein are structurally aligned and coloured in light blue. **a**. The allosteric site surface (grey) bound to S41 (red) of GK is shown, and residues 93 to 99 (blue) are missing in the 3S41 structure. **b**. The binding site surface (grey) bound to the triclosan analog (red) of PfENR is shown, with residues 318 to 368 (blue) missing in the 2OP1 structure.

Another interesting example of ghost interactions involves the case of *Plasmodium falciparum* enoyl acyl carrier protein reductase (PfENR), a drug target to treat malaria. The flexibility of the active site suggests it could be optimised to improve compound potency against Plasmodium parasites. The crystal structure of PfENR bound to triclosan analog (inhibitor, PDB: 2OP1) reveals several interactions in the active site^41^ (Figure 7b). The AF2 structure completes the region from R318 to F368, exhibiting ghost interactions between the inhibitor in the active site and residues A319, A320, I323 and F368 through their hydrophobic moieties and pi-pi interactions closing the active site.

## Discussion

Extensive studies about how disordered regions modulate ligand binding have shown a continuum between extreme cases such as pre-formed (ordered) or completely disordered binding sites, depending on each particular intrinsically disordered protein ^42,43^. Whatever the ligand-binding approach a disordered region could show, these interactions with ligands are the essential requirements to sustain the biological function. These interactions broaden the ligand binding capacity due to the presence of flexible regions that can fit ligands with very different sizes combining several low strength interactions^42^. The study of these interactions between disordered regions and ligands is difficult since contact information is unavailable in crystallographic structures, in contrast with the study of ordered binding sites ^6,44^. In this work we have shown that AF2 can estimate the presence of interactions between ligands and residues belonging to disordered regions, which we called “ghost interactions” as they are missing in the crystallography derived structures. First, we found that AF2 models are good predictors of regions with high flexibility or the presence of disorder. We tested this capacity on order-disorder regions (different conformers showing a given region as ordered and disordered), which allowed us to estimate the AF2 error rate. We found that AF2 predicted ODR regions with an error in the same order as the conformational diversity of these regions (Figure 2). The error for ODR lengths between 5 and 100 residues long, lies in the RMSD interval between 1.1 and 1.6 Å (Figure 3). Using holo forms from ∼11000 proteins, we found that the relative error in AF2 estimation of contacts with ordered regions has a median of ∼8%. Selecting those holo forms with missing regions near the binding region (1113 structures) we were able to characterise the presence of ghost interactions. These regions appearing as missing in the PDB structure are modelled by AF2. Residues making ghost interactions are mostly buried and differentially evolutionary conserved, both in reference to the rest of the residues in the same ODR region and to the rest of the protein (Figure 5). As disordered regions generally have poor evolutionary conservation ^45^, the fact that residues with ghost interactions are evolutionary conserved can indicate their biological relevance in terms of protein-ligand interactions. In fact, we showed the uniqueness of these residues both in their degree of conservation, amino acid composition and RSA in reference to other disordered residues. Taking advantage of these defining features for the detection of ghost interactions with the combined use of crystallographic structures and AF2 models could help to understand the contribution of disorder in the stabilisation of ligands, explain differentially conserved residues observed in disordered regions, and provide insights into the transit of ligands to the binding site. In this sense, our findings also support the current tendency to explore intrinsically disordered proteins as potentially valuable targets for drug development, given their biological relevance and role in associated diseases.

## Acknowledgements

MSF, NP, TS, LRS, SFA and GP are researchers and NE and JMD are PhD fellows from Consejo Nacional de Investigaciones Científicas y Técnicas (CONICET). This work was supported by Universidad Nacional de Quilmes (PUNQ 1004/11), Agencia Nacional de Promoción Científica y Tecnológica (PICT-2014-3430) and the European Union’s Horizon 2020 Research and Innovation Programme (Grant Agreement N° 778247 and N° 823886).

## Author contributions

N.E. provided CoDNaS and MobiDB conformers and other datasets, prepared example visualisations. T.S. worked on calculations and structural comparisons. J.M.D. mapped the PDB-UniProt, and worked on the statistics and visualisation. L.R.S. selected and described biological examples and worked with visualisation. N.P. assisted in conceptualization and manuscript revision. S.F.A. and M.S.F provided funding and conceptualization. G.P conceptualised and designed the study. All authors contributed to the manuscript writing.

## Methods

### Dataset and order-disorder regions

ODRs represent missing regions in at least one structure (conformer) with the same region being structured (ordered) in at least another one. Conformer information was extracted from the CoDNaS database ^32^. Order-disorder transition regions (ODRs) were defined using disorder information from MobiDB ^33^ mapping the structures to their Uniprot accession to identify individual proteins. We only considered proteins with at least one ODR with more than 5 and up to 100 consecutive residues. For each of these proteins we downloaded the AF2 model from the EMBL-EBI AlphaFold Protein Structure Database at https://ftp.ebi.ac.uk/pub/databases/alphafold. Initially, we found 170,331 disordered fragments, 57,364 ordered conformers, and 24,380 AF2 models. After filtering structures by missing length and removing the structures that have disorder regions near the starts or ends of the protein, we mapped each of the corresponding AF2 (UniProt) positions to the corresponding PDB positions to allow comparison of the each same residue in the PDB and the AF2 models. This was done using the PDBSWS mapping resource found in http://bioinf.org.uk/servers/pdbsws/ ^46^. After this, we also filtered the proteins that had PDB - UniProt mapping with errors. Our final dataset was composed of 10,908 fragments in 4,097 protein structures.

### Comparison of structures and fragments

The AF2 models and corresponding ODRs were structurally aligned using the R package Bio3D ^47^. We quantified the structural similarity of each fragment using the alpha carbon-Root Mean Square Deviation (C⍺-RMSD). When more than two conformers showing the ODR as ordered were found, the fragment with the lowest C⍺-RMSD was registered. For the bootstrapping analysis, we randomly selected fragments of consecutive residues with the same size as the missing fragment from other regions in the same protein, and aligned each fragment against the equivalent ODR of the AF2 model. We generated 100 alignments with alternative fragments for each conformer and the C⍺-RMSD was calculated between each alignment pair. To estimate the conformational diversity of the ODR regions with more than one conformer showing that region as ordered, we calculated the C⍺-RMSD between each fragment and registered the maximum value per ODR.

### Using AF2 to predict contacts

Using CoDNaS conformers and the BioLiP database, we obtained 11,113 PDB files for protein holo forms. To determine AF2 contacts, we aligned AF2 models with the holo forms and then transplanted ligands to the AF2 file. Following previous validation of transplanted ligands ^34^ in AF2 models, we estimated the “transplant clash score” (TCS) defined as:

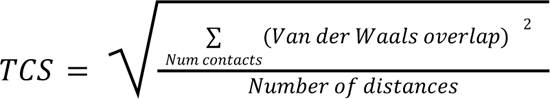

To further validate the models, we compared the distribution of clashes with the corresponding holo forms. Contacts between ligands and PDB files and AF2 models were calculated with the R package Bio3D. Ligands that made contact with more than 70 residues were considered outliers. We also removed the ligands that are peptides because they have noise on the relevant contacts detection. To remove them we look for those containing the word “peptide” within the PDB download service tag “chem_comp” and subtag “type” (see https://data.rcsb.org/redoc/index.html for more information). To facilitate the study we used only A chains with only the first ligand reported. All pairs of structures with RMSD larger than 30Å were also removed as they were more than likely mistakes.

### Ghost interaction analysis

A “ghost interaction” involves residues in the AF2 model that are located in a missing region, and that make noncovalent contacts (at distance equal or less than 5 Å) with the transplanted ligand. Using the reported missing regions (disordered regions) derived from MobiDB database in the holo forms aligned with their corresponding AF2 models, as mentioned above, we obtained 456 proteins with 1,113 PDB structures where the ligands were making contacts with AF2 models, thus showing “ghost interactions”. The amino acid distributions were calculated with the Composition Profiler Tool server http://www.cprofiler.org/cgi-bin/profiler.cgi ^48^.

### Conservation and Relative Surface Area

For each protein with “ghost interactions” we ran BLAST sequence similarity searches with default parameters to retrieve sequences with more than 30% identity and E-value> 1.10^-10^. Redundancy was reduced using CD-Hit ^49^ by removing sequences with more than 90% identity. The remaining sequences were aligned with the Muscle program ^50^. Conservation was calculated with the AL2CO software ^51^, using unweighted variance measure as the main parameters. The other 8 available parameters were highly correlated for our dataset.

Accessible Surface Area was calculated using the Define Secondary Structure of Proteins software ^52^. The raw values of the Accessible Surface Area were divided by the free residue surface for the corresponding residue, in order to obtain the Relative Solvent Accessible Area (RSA).

## Data availability

Code and datasets are available at: https://gitlab.com/sbgunq/publications/ghost_interactions_2023/

## Supplementary information

### Supplementary Figures

**Supplementary Figure 1:**
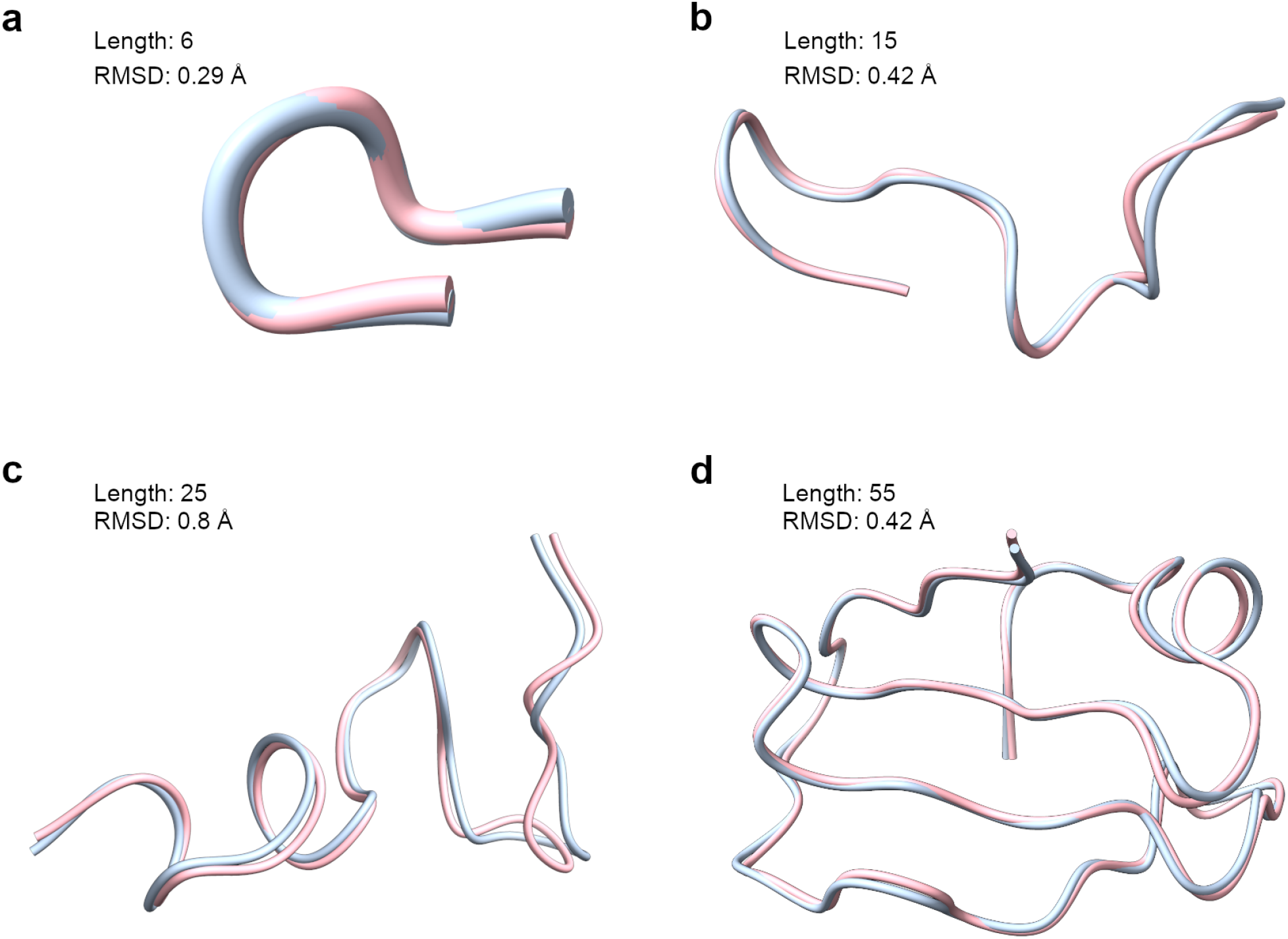
Some examples of AF2 capacity to predict order-disorder regions. In light blue AF2 structures, in light pink experimental structures. **a.** PDB= 1MEW_A with ODR= 6 residues, **b.** PDB= 5MNI_B with ODR= 15, **c.** PDB= 1S5P_A with ODR= 25, **d.** PDB= 4IMA_C with ODR= 55.

**Supplementary Figure 2:**
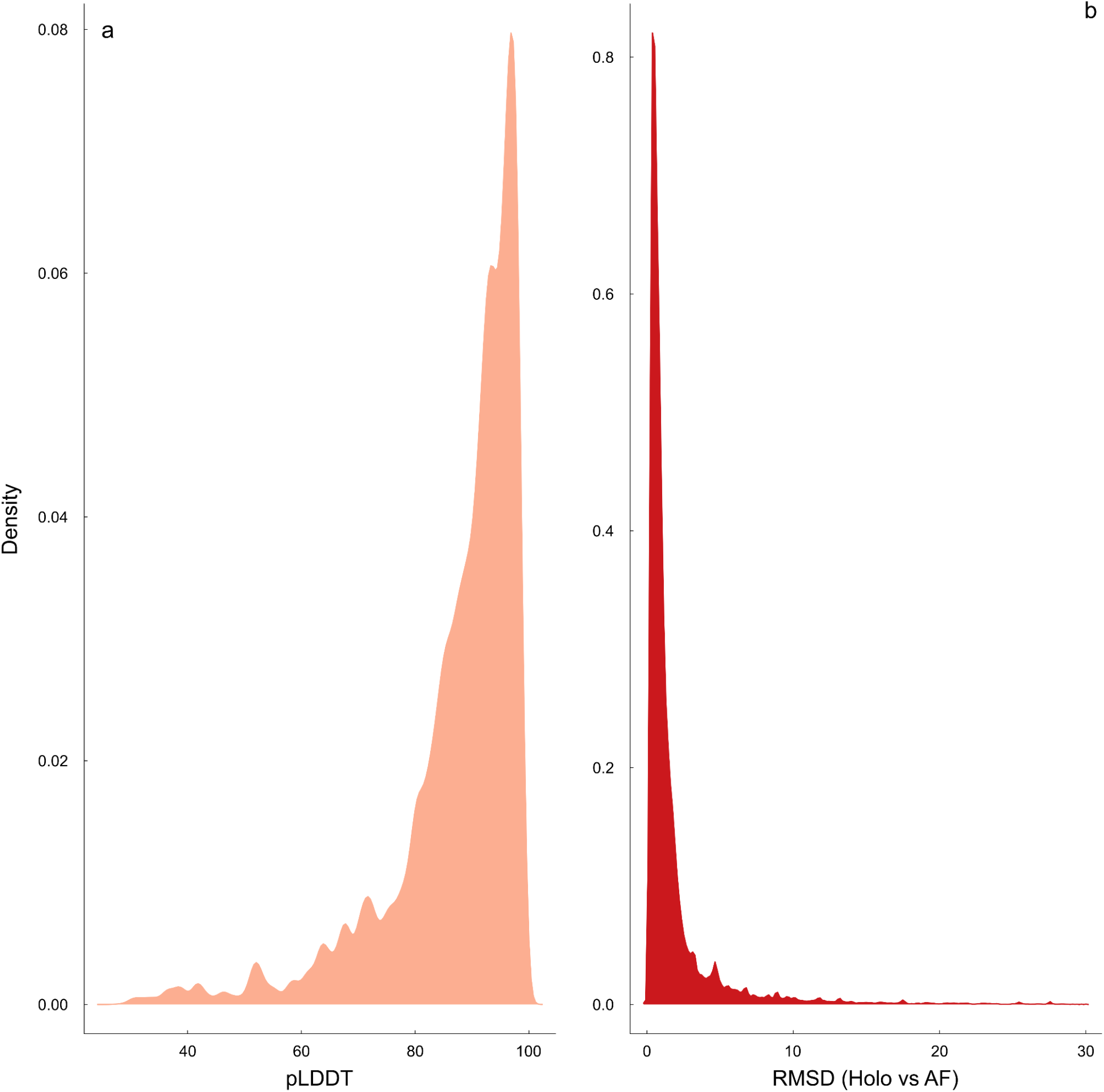
pLDDT and RMSD (holo vs AF2 models) distributions for 11323 proteins with cognate ligands (holo forms). For each of these proteins we obtained the corresponding AF2 models. The distribution of their pLDDT is shown in the left panel (average of 87.4). In the right panel we show the global RMSD obtained between each protein and its corresponding AF2 model (Average 1.91). Both distributions show the excellent quality of the AF2 models used.

**Supplementary Figure 3:**
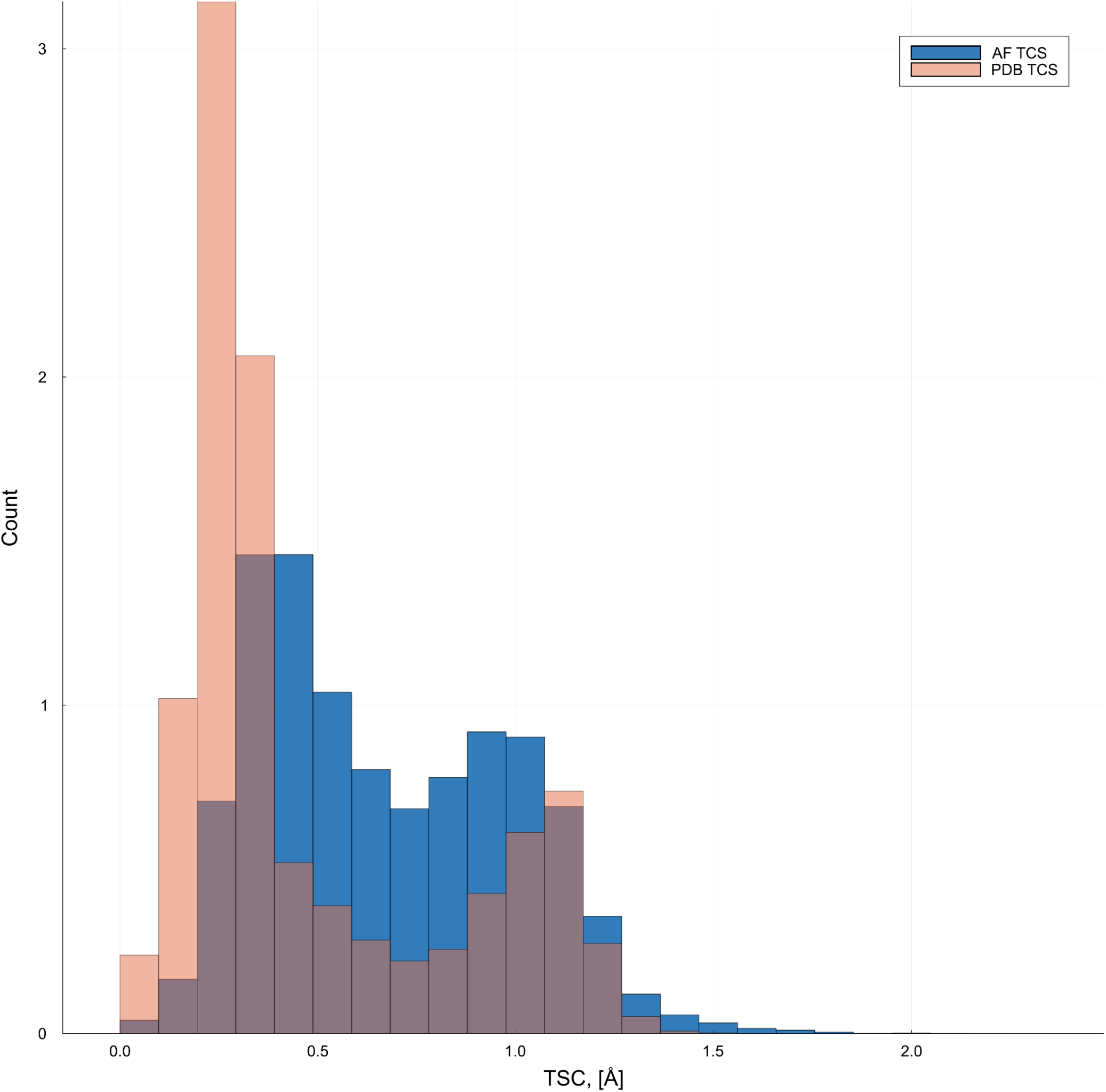
Clash distribution between ligand and protein for PDB structures and AF2 models. Following Hekkelman *et al* ^34^ we estimated the proportion of clashes between ligands in the holo form of protein structures derived from PDB and derived from AF2 using a score called the “transplant clash score” (TCS) (See Methods in the Main Manuscript). Ligand positions in AF2 were derived after structural alignment with the protein holo form derived from PDB. The figure shows that both distributions are similar, indicating that our AF2-ligand structures have the same error distribution as the one derived from PDB. The 90 percentile of TSC for holo forms from PDB is 1.07 Å and for the AF2 model is 1.10 Å.

**Supplementary Figure 4:**
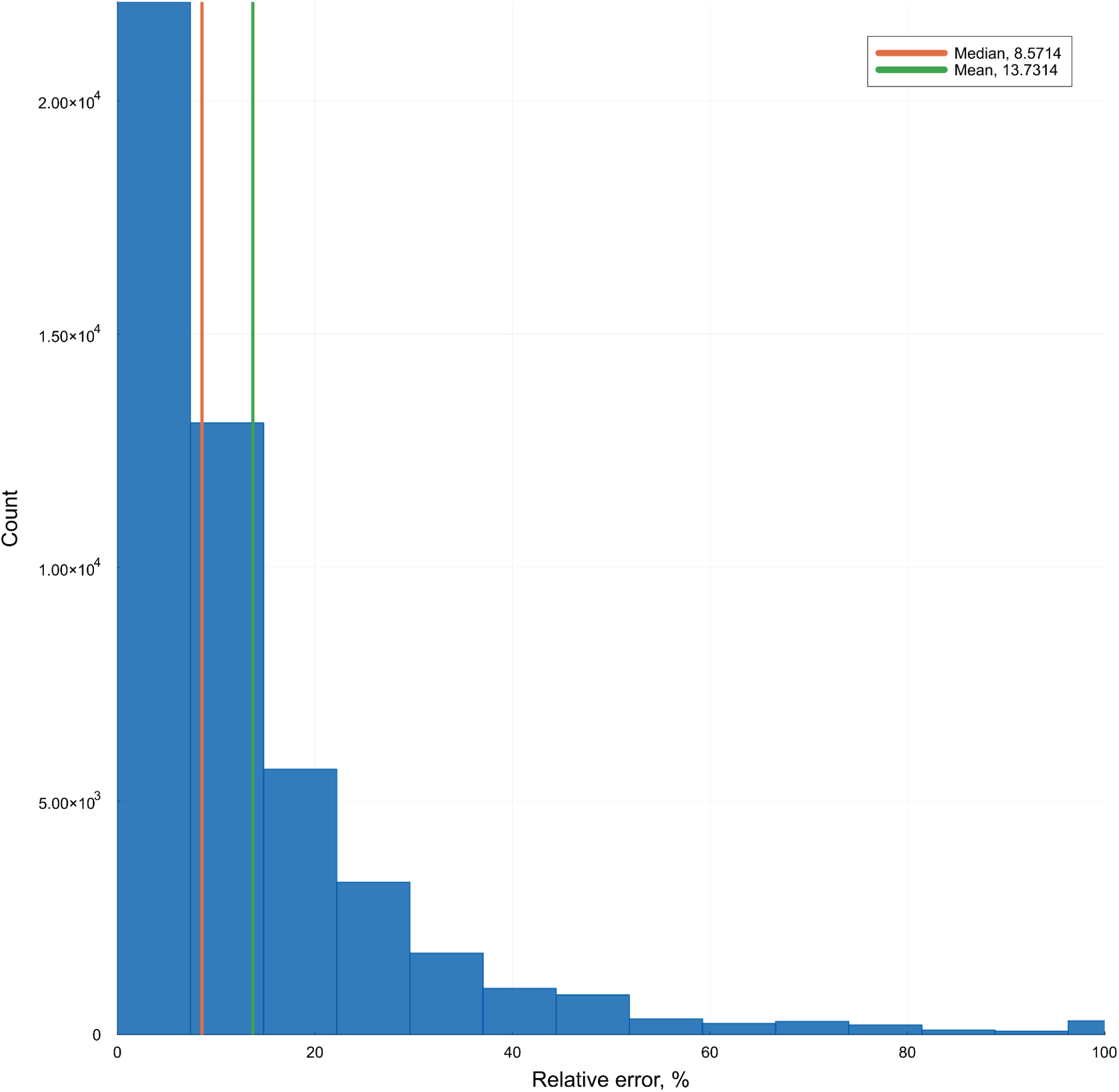
Relative error in contacts prediction. Distribution of the contact difference between PDB contacts and AF2 predicted contacts, normalised by the number of PDB contacts (relative error). The median of this distribution is 8.57.

**Supplementary Figure 5:**
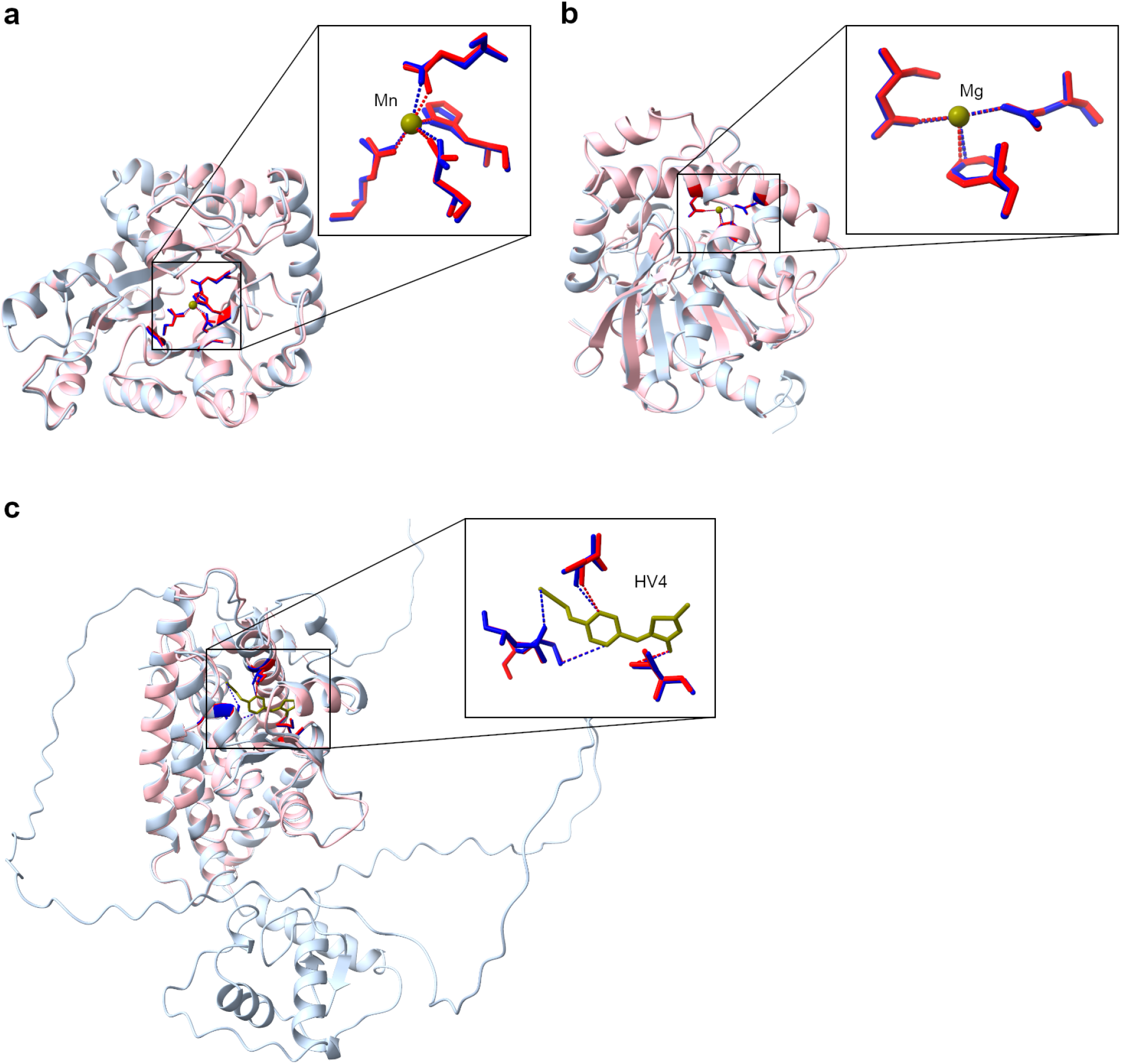
Example of how AF2 predicts contacts in ordered regions. For all panels AF2 models are shown in light blue and experimental structures in light pink. Biological ligands in green, AF2 model residues that make contacts in blue, and the same PDB residues in red. **a.** 2UQL_A and AF2 structure for O50580. Non optimised RMSD is 0.23 Å, pLDDT = 98.74, both have 7 contacts with Mn. **b.** 1Y37_A and AF2 Q1JU72, RMSD= 1.02 Å, pLDDT = 98.72, both present 9 contacts with MG. **c.** 6E5A_A and AF2 P37231, RMSD= 0.68 Å, pLDDT = 70.48, the PDB makes 15 contacts and AF2 makes 17 contacts with HV4. The real number of interactions are reduced for simplicity.

**Supplementary Figure 6:**
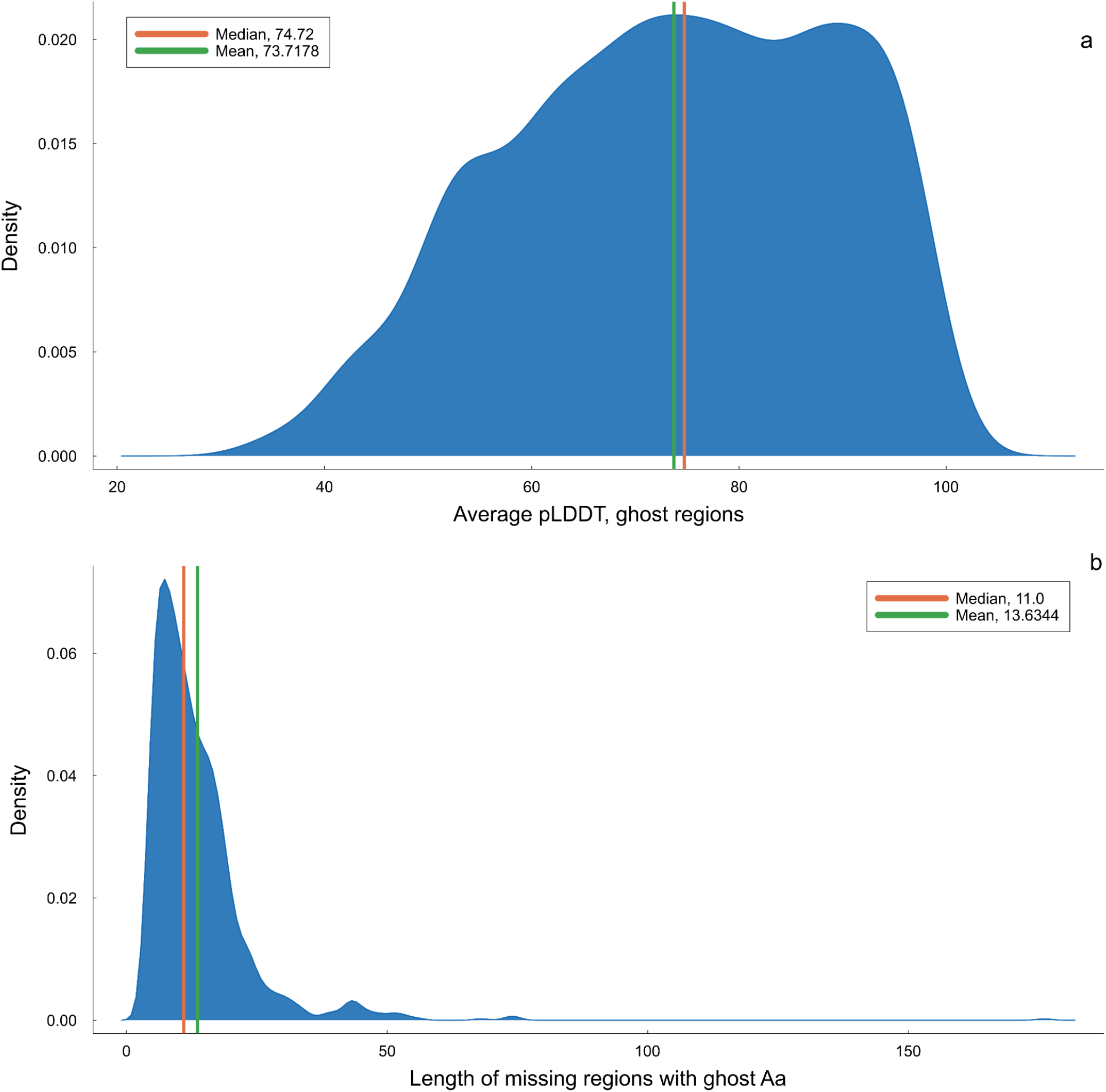
pLDDT and Length distributions for holo forms containing regions with ghost interactions.

**Supplementary Figure 7:**
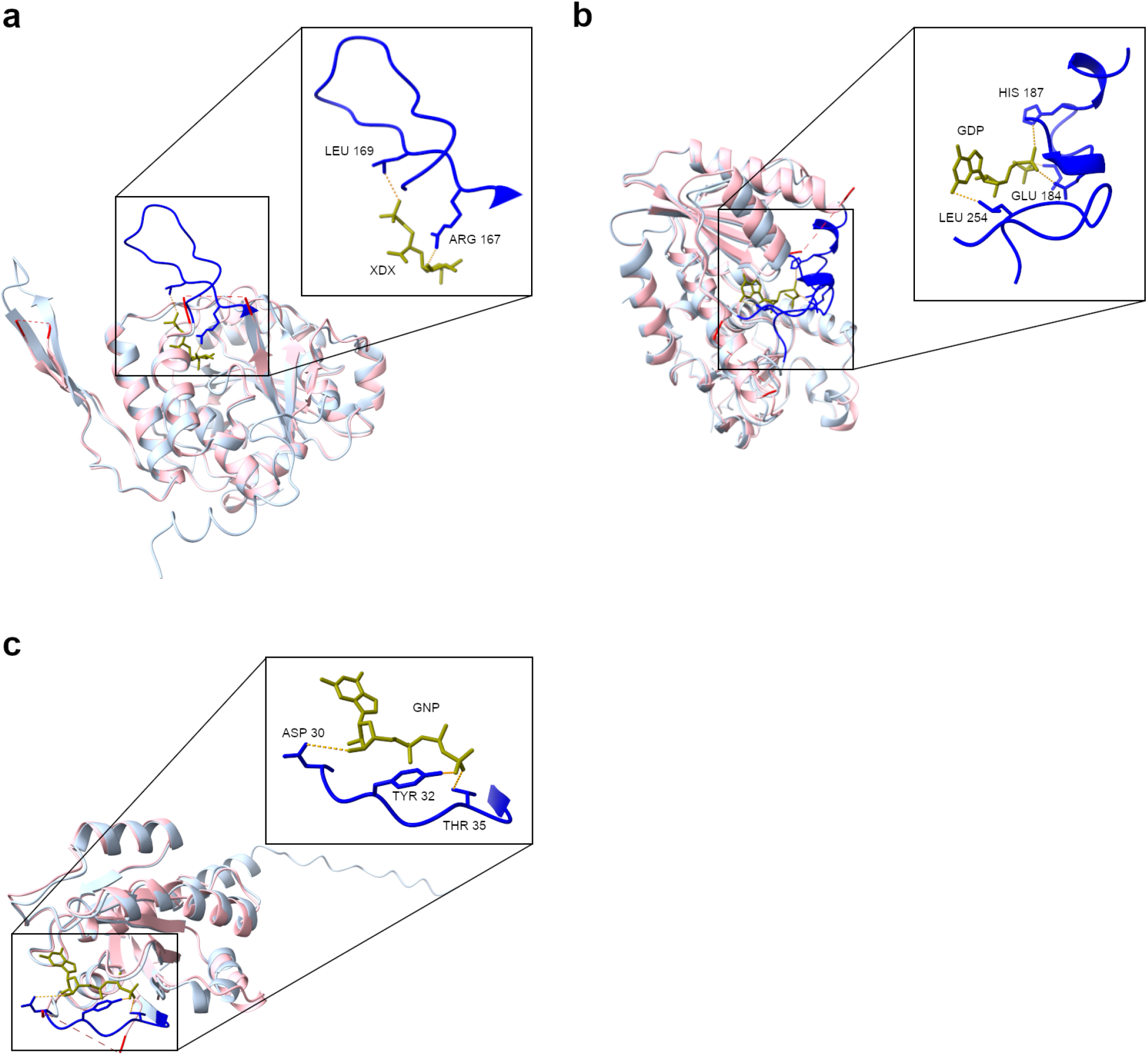
Ghost interactions and ODR lengths. For all panels AF2 models are shown in light blue and experimental structures in light pink. Missing regions in experimental structures are shown as red dashes, AF2 models of these regions are shown in blue. The real number of ghost interactions are reduced for simplicity. **a.** 5JUC_A and AF2 P9WMW9, RMSD= 0.83 Å, pLDDT = 87.45, 4 ghost interactions with ligand XDX (ARG167, PRO168, LEU169, GLY182). **b.** 3SIX_A and AF2 Q9AQ17, RMSD= 1.46 Å, pLDDT = 92.18, 8 ghost interactions with GDP (GLY179, GLU182, ILE184, MET185, HIS187, PRO253, LEU254, HIS255). **c.** 5WLB_A and AF P01116, RMSD= 1.52 Å, pLDDT = 92.69, 5 ghost interactions with GNP (GLU31, TYR32, ASP33, PRO34, THR35).

**Supplementary Figure 8:**
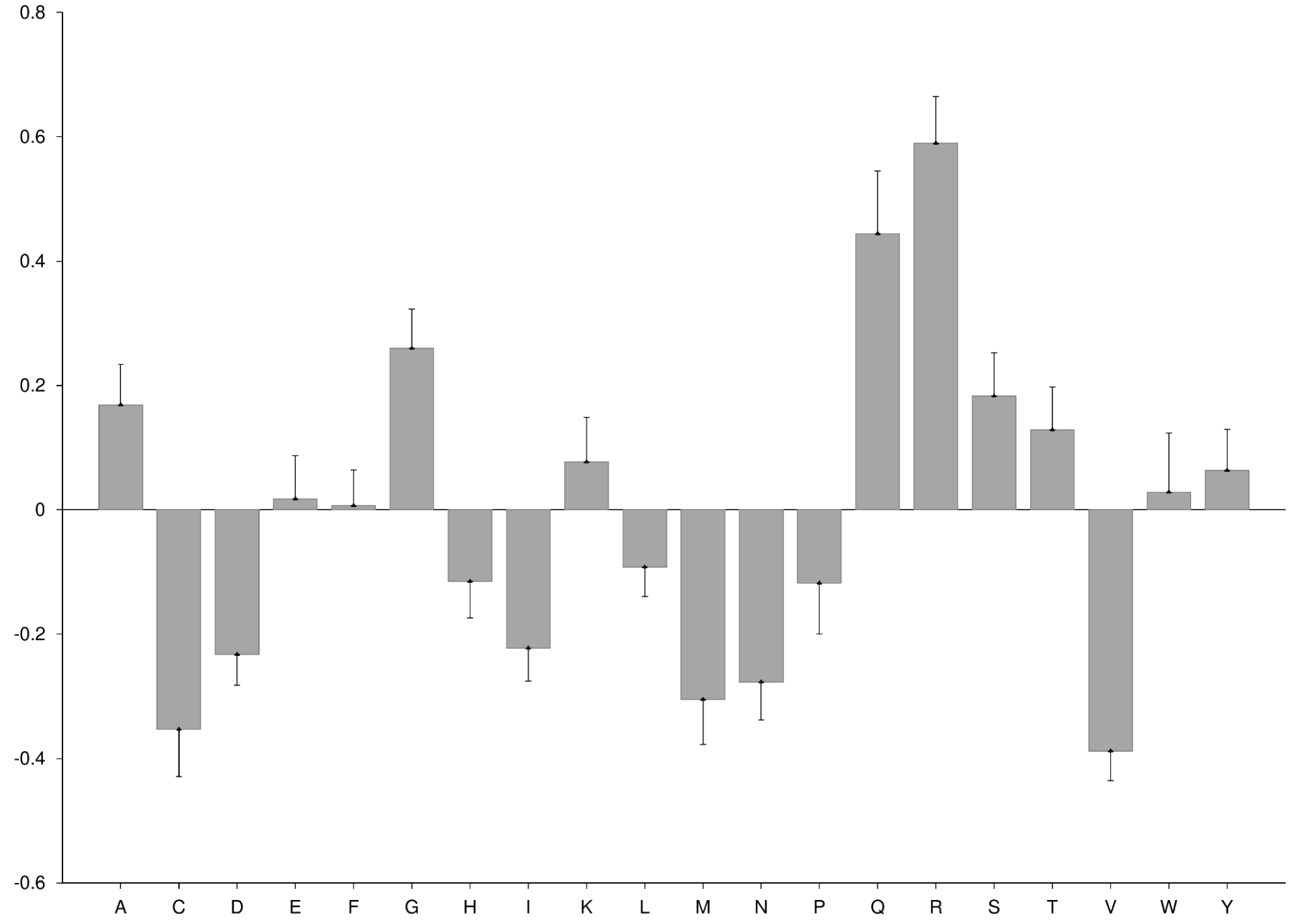
Amino acid distribution of residues with ghost interactions taking as background distributions those residues making contacts with ligands in ordered regions of the proteins.

**Supplementary Figure 9:**
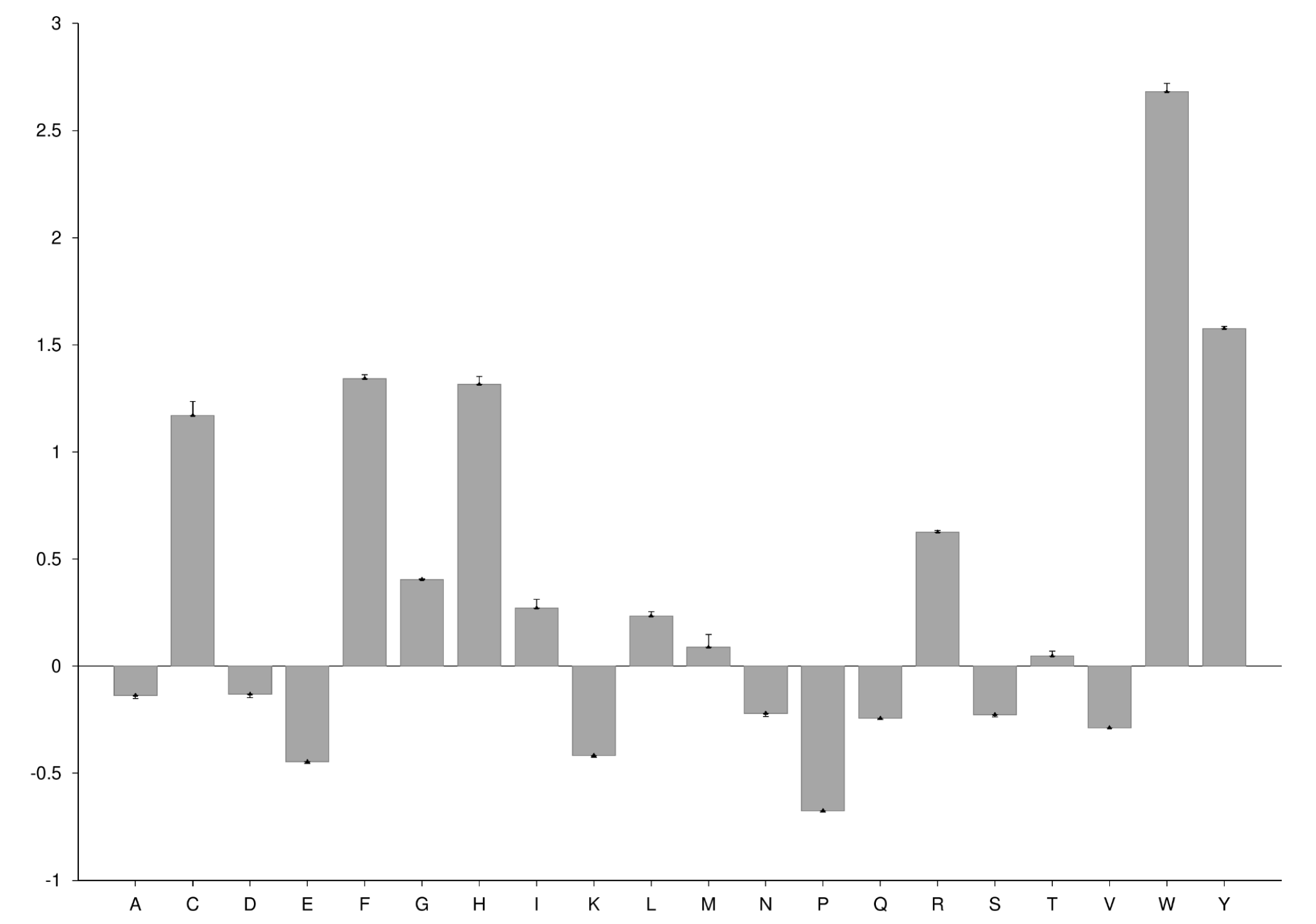
Amino acid distribution of residues with ghost interactions taking as background distributions Disprot 3.4 reference distribution.

### Supplementary Tables

**Supplementary Table 1:** Wilcoxon rank-sum test p-values for figure 2.

| RMSD 1 | RMSD 2 | p-value |
| --- | --- | --- |
| All Fragments | pLDDT < 70 | < 0.001 |
| All Fragments | pLDDT >= 70 | < 0.001 |
| All Fragments | 1 conformer | < 0.001 |
| All Fragments | 2 or more conformers | < 0.001 |
| All Fragments | Bootstrapped fragments | < 0.001 |
| All Fragments | Disorder regions | < 0.001 |
| pLDDT < 70 | pLDDT >= 70 | < 0.001 |
| pLDDT < 70 | 1 conformer | < 0.001 |
| pLDDT < 70 | 2 or more conformers | < 0.001 |
| pLDDT < 70 | Bootstrapped fragments | < 0.001 |
| pLDDT < 70 | Disorder regions | < 0.001 |
| pLDDT >= 70 | 1 conformer | < 0.001 |
| pLDDT >= 70 | 2 or more conformers | < 0.001 |
| pLDDT >= 70 | Bootstrapped fragments | < 0.001 |
| pLDDT >= 70 | Disorder regions | < 0.001 |
| 1 conformer | 2 or more conformers | < 0.001 |
| 1 conformer | Bootstrapped fragments | < 0.001 |
| 1 conformer | Disorder regions | < 0.001 |
| 2 or more conformers | Bootstrapped fragments | < 0.001 |
| 2 or more conformers | Disorder regions | < 1 |
| Bootstrapped fragments | Disorder regions | < 0.001 |

p-values Table 2 (Figure 3, Wilcoxon rank-sum test).

**Supplementary Table 2:**
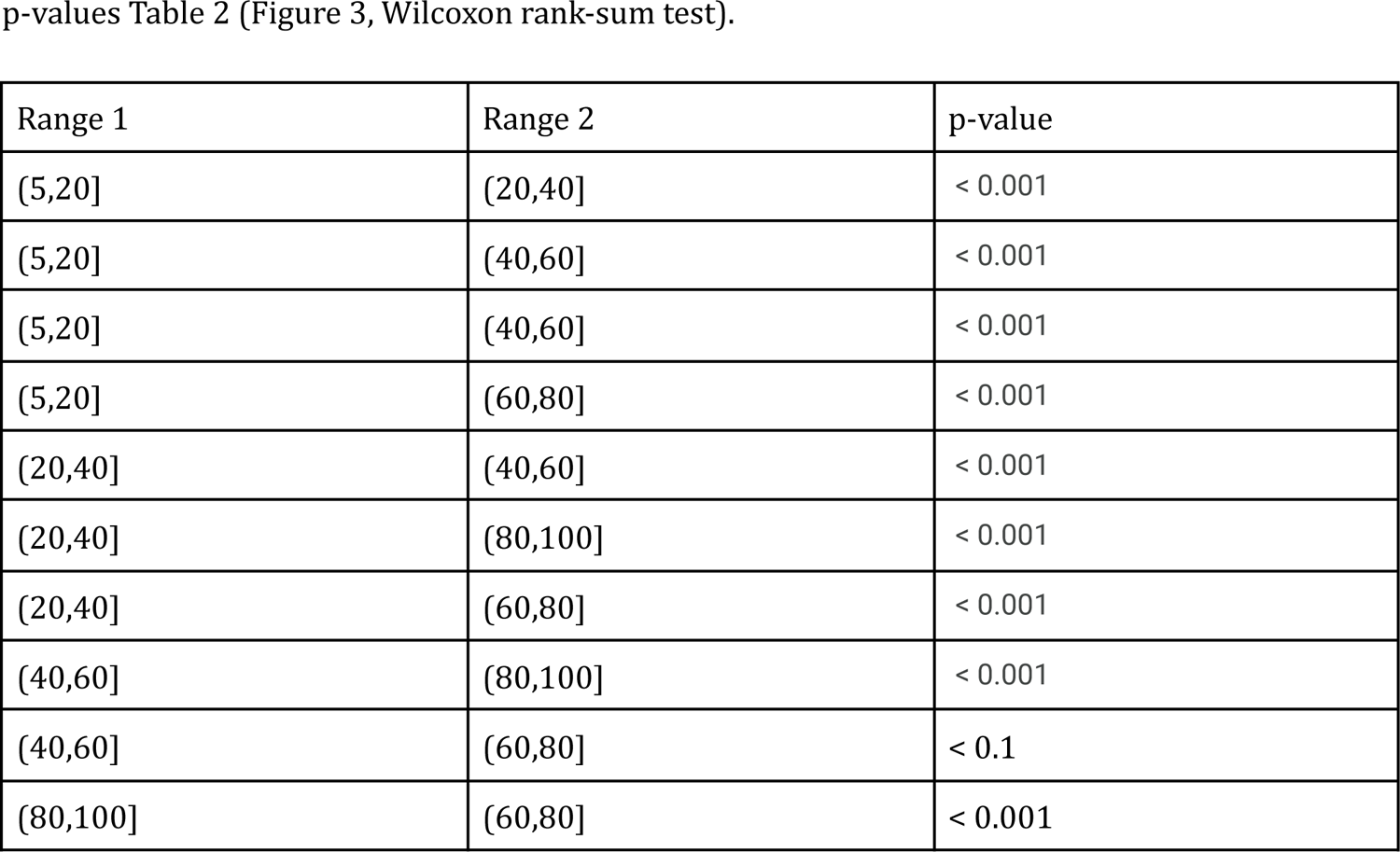
Wilcoxon rank-sum test p-values for figure 3.

**Supplementary Table 3:** Wilcoxon rank-sum test p-values for figure 5.

Conservation
| Group 1 | Group 2 | p-value |
| --- | --- | --- |
| Ghost_cons | AF_contacts_cons | < 0.001 |
| Ghost_cons | Missing_no_ghost_cons | < 0.001 |
| AF_contacts_cons | Missing_no_ghost_cons | < 0.001 |

RSA
| Group 1 | Group 2 | p-value |
| --- | --- | --- |
| Ghost_RSA | AF_contacts_RSA | < 0.001 |
| Ghost_RSA | missings_RSA_no_ghost | < 0.001 |
| AF_contacts_RSA | missings_RSA_no_ghost | < 0.001 |

pLDDT
| Group 1 | Group 2 | p-value |
| --- | --- | --- |
| Ghost_pLDDT | missings_pLDDT | < 0.001 |
| Ghost_pLDDT | AF_contacts_pLDDT | < 0.001 |
| missings_pLDDT | AF_contacts_pLDDT | < 0.001 |

**Supplementary Table 4:**
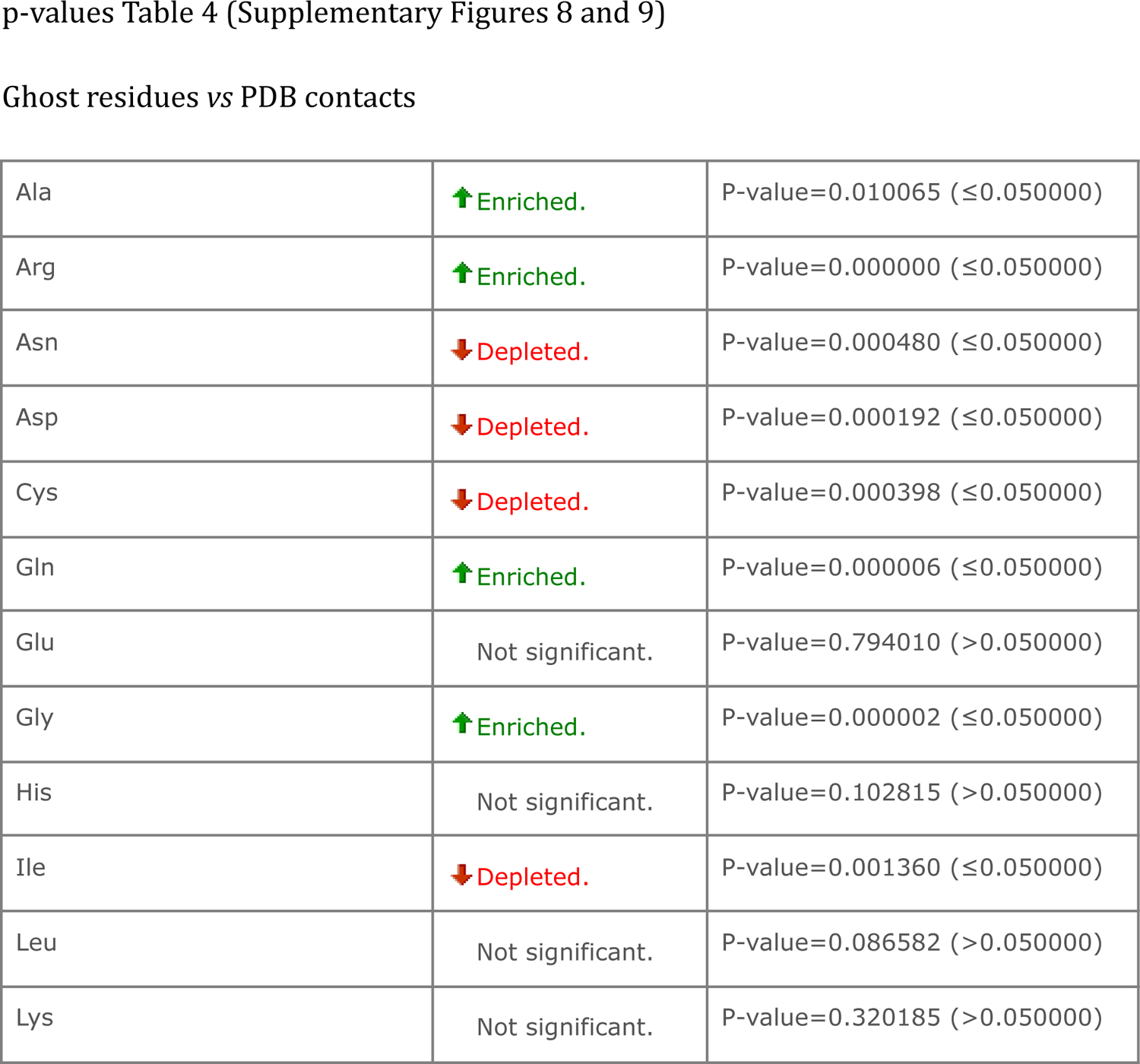
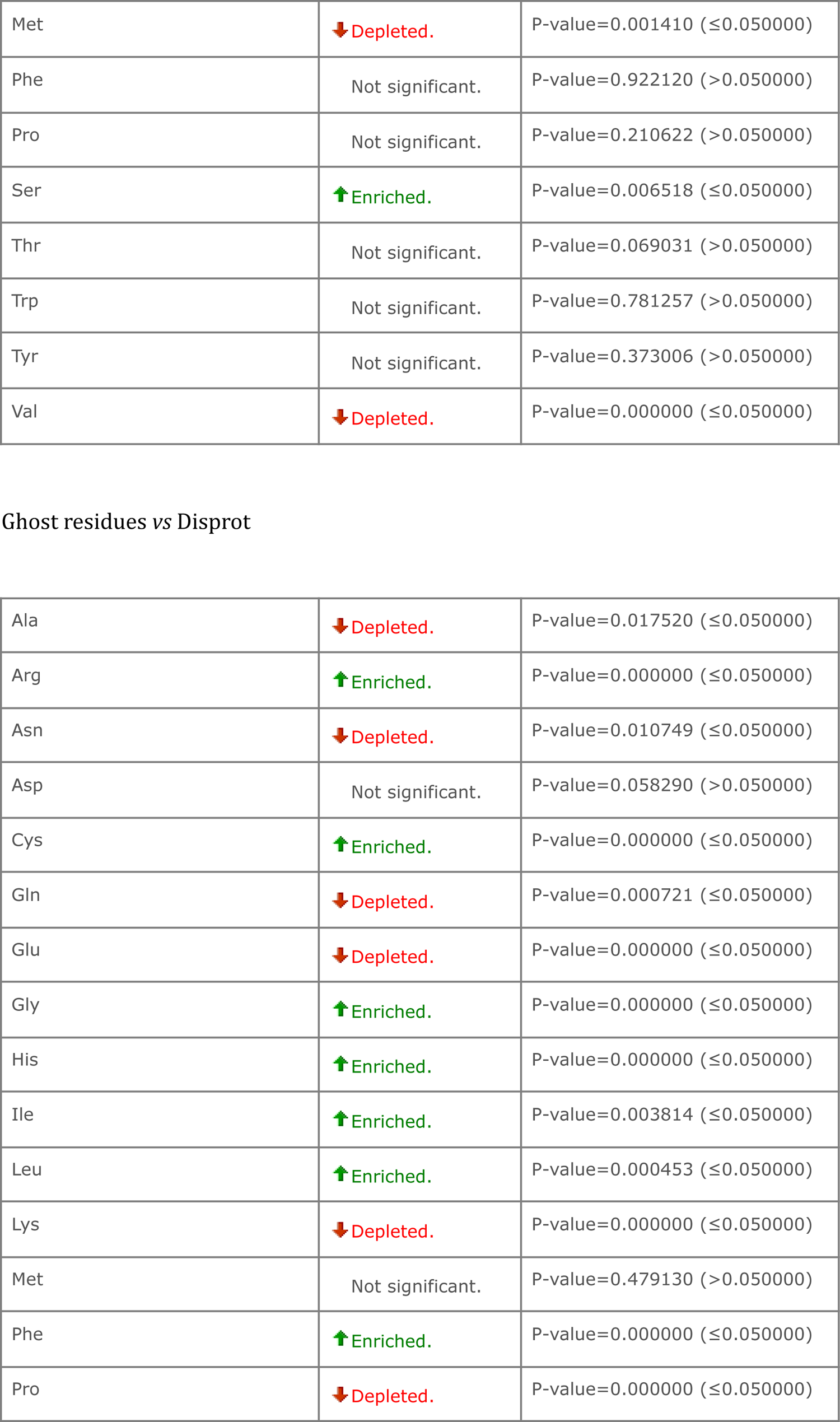
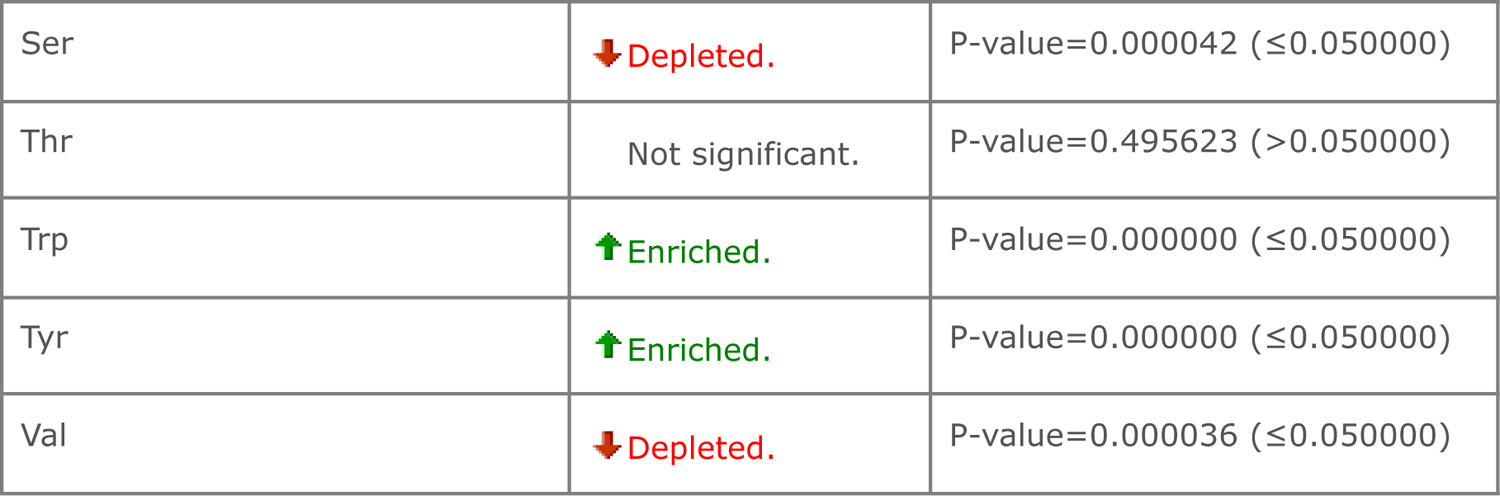
p-values for enrichment and depletion of ghost residues.

Ghost residues vs PDB contacts
|  |  |  |
| --- | --- | --- |
| Ala | ↑ Enriched. | P-value=0.010065 ($\leq 0.050000$) |
| Arg | ↑ Enriched. | P-value=0.000000 ($\leq 0.050000$) |
| Asn | ↓ Depleted. | P-value=0.000480 ($\leq 0.050000$) |
| Asp | ↓ Depleted. | P-value=0.000192 ($\leq 0.050000$) |
| Cys | ↓ Depleted. | P-value=0.000398 ($\leq 0.050000$) |
| Gln | ↑ Enriched. | P-value=0.000006 ($\leq 0.050000$) |
| Glu | Not significant. | P-value=0.794010 ($> 0.050000$) |
| Gly | ↑ Enriched. | P-value=0.000002 ($\leq 0.050000$) |
| His | Not significant. | P-value=0.102815 ($> 0.050000$) |
| Ile | ↓ Depleted. | P-value=0.001360 ($\leq 0.050000$) |
| Leu | Not significant. | P-value=0.086582 ($> 0.050000$) |
| Lys | Not significant. | P-value=0.320185 ($> 0.050000$) |
| Met | ↓ Depleted. | P-value=0.001410 ($\leq 0.050000$) |
| Phe | Not significant. | P-value=0.922120 ($> 0.050000$) |
| Pro | Not significant. | P-value=0.210622 ($> 0.050000$) |
| Ser | ↑ Enriched. | P-value=0.006518 ($\leq 0.050000$) |
| Thr | Not significant. | P-value=0.069031 ($> 0.050000$) |
| Trp | Not significant. | P-value=0.781257 ($> 0.050000$) |
| Tyr | Not significant. | P-value=0.373006 ($> 0.050000$) |
| Val | ↓ Depleted. | P-value=0.000000 ($\leq 0.050000$) |
| Ghost residues vs Disprot |  |  |
| Ala | ↓ Depleted. | P-value=0.017520 ($\leq 0.050000$) |
| Arg | ↑ Enriched. | P-value=0.000000 ($\leq 0.050000$) |
| Asn | ↓ Depleted. | P-value=0.010749 ($\leq 0.050000$) |
| Asp | Not significant. | P-value=0.058290 ($> 0.050000$) |
| Cys | ↑ Enriched. | P-value=0.000000 ($\leq 0.050000$) |
| Gln | ↓ Depleted. | P-value=0.000721 ($\leq 0.050000$) |
| Glu | ↓ Depleted. | P-value=0.000000 ($\leq 0.050000$) |
| Gly | ↑ Enriched. | P-value=0.000000 ($\leq 0.050000$) |
| His | ↑ Enriched. | P-value=0.000000 ($\leq 0.050000$) |
| Ile | ↑ Enriched. | P-value=0.003814 ($\leq 0.050000$) |
| Leu | ↑ Enriched. | P-value=0.000453 ($\leq 0.050000$) |
| Lys | ↓ Depleted. | P-value=0.000000 ($\leq 0.050000$) |
| Met | Not significant. | P-value=0.479130 ($> 0.050000$) |
| Phe | ↑ Enriched. | P-value=0.000000 ($\leq 0.050000$) |
| Pro | ↓ Depleted. | P-value=0.000000 ($\leq 0.050000$) |
| Ser | ↓ Depleted. | P-value=0.000042 ($\leq 0.050000$) |
| Thr | Not significant. | P-value=0.495623 ($> 0.050000$) |
| Trp | ↑ Enriched. | P-value=0.000000 ($\leq 0.050000$) |
| Tyr | ↑ Enriched. | P-value=0.000000 ($\leq 0.050000$) |
| Val | ↓ Depleted. | P-value=0.000036 ($\leq 0.050000$) |

